# Oligomeric α-synuclein-specific degradation by HtrA2/Omi to bestow a neuroprotective function

**DOI:** 10.1101/468967

**Authors:** Hea-Jong Chung, Mohammad Abu Hena Mostofa Jamal, Mashiar Rahman, Hyeon-Jin Kim, Seong-Tshool Hong

**Affiliations:** Department of Biomedical Sciences and Institute for Medical Science, Chonbuk National University Medical School, Jeonju, Chonbuk 54907, South Korea; JINIS BDRD institute, JINIS Biopharmaceuticals Co., 913 Gwahak-Ro, Bongdong, Wanju, Chonbuk 55321, South Korea; Department of Biotechnology and Genetic Engineering, Islamic University, Kushtia-7003, Bangladesh

## Abstract

Although the malfunction of HtrA2/Omi leads to Parkinson’s disease (PD), the underlying mechanism has remained unknown. Here, we showed that HtrA2/Omi specifically removed oligomeric α-Syn but not monomeric α-Syn to protect oligomeric α-Syn-induced neurodegeneration. Experiments using mnd2 mice indicated that HtrA2/Omi degraded oligomeric α-Syn specifically without affecting monomers. Transgenic *Drosophila melanogaster* experiments of the co-expression α-Syn and HtrA2/Omi and expression of genes individually also confirmed that pan-neuronal expression of HtrA2/Omi completely rescued Parkinsonism in the α-Syn-induced PD *Drosophila* model by specifically removing oligomeric α-Syn. HtrA2/Omi maintained the health and integrity of the brain and extended the life span of transgenic flies. Because HtrA2/Omi specifically degraded oligomeric α-Syn, co-expression of HtrA2/Omi and α-Syn in *Drosophila* eye maintained a healthy retina, while the expression of α-Syn induced retinal degeneration. This work showed that the bacterial function of HtrA to degrade toxic misfolded proteins is evolutionarily conserved in mammalian brains as HtrA2/Omi.

## INTRODUCTION

Parkinson’s disease (PD) is one of the most common neurodegenerative diseases, characterized by the progressive loss of dopaminergic neurons in the substantia nigra of the central nervous system^1^. The degeneration of the dopaminergic neurons of the substantia nigra results in clinical manifestations such as motor impairments, which involve resting tremor, bradykinesia, postural instability, gait difficulty and rigidity^2^. Although the degeneration of dopaminergic neurons directly leads to the clinical manifestations of PD, the pathogenic mechanism underlying the degeneration of dopaminergic neurons at the molecular level is still unclear. The most manifested pathophysiological feature of dopaminergic neurons of PD is an abnormal accumulation of oligomeric α-Synuclein (α-Syn) in the form of Lewy bodies and Lewy neuritis inside neurons, which represent the major hallmarks of PD^3^. α-Syn is a 140-a.a. presynaptic protein that plays an important role in maintaining a supply of synaptic vesicles in presynaptic terminals^4, 5^. α-Syn performs its normal biological function in neurons if present as a monomer. However, the monomeric form of α-Syn is naturally prone to adopt a β-sheet conformation to form oligomeric aggregates^6^. The oligomeric α-Syn has very strong neurotoxicity such that the aggregation plays a causative role in dopaminergic neuronal degeneration^7, 8^.

Since the accumulation of misfolded α-Syn is key to the pathology of PD, the question of how misfolded α-Syn is degraded by neurons has been actively investigated. Investigations over the last decades have elucidated that the ubiquitin-proteasome system (UPS) and the autophagy-lysosomal pathway (ALP) work in conjunction to degrade α-Syn^9, 10^. However, neither UPS nor ALP are specific pathways for α-Syn degradation but rather are for general intracellular protein turn-over pathways^11-13^. More importantly, none of the pathways have shown selectivity toward oligomeric α-Syn, and thus almost all the wasted proteins in cells, including monomeric α-Syn, are degraded by the pathways. Therefore, the mechanisms for oligomeric α-Syn degradation pathway to relieve the toxicity of oligomeric α-Syn in neurons remain completely unknown.

One of the dilemmas in neurons concerns the lack of toxicity of native monomeric α-Syn; rather, it is essential for proper neuronal functions, while oligomeric α-Syn is very neurotoxic^14^. Because monomeric α-Syn plays indispensable roles in neurons, the α-Syn knock-out mouse shows impaired spatial learning and working memory^15^. Considering that monomeric α-Syn has a naturally strong tendency to self-aggregate into neurotoxic oligomers^16, 17^, it is reasonable to speculate that neurons have an unknown pathway that specifically recognizes oligomeric α-Syn only to degrade oligomeric α-Syn without affecting the monomeric form.

HtrA2/Omi is a homolog of the bacterial heat shock protein HtrA (also known as DegP), which protects bacteria at elevated temperatures by specifically recognizing denatured proteins to degrade those proteins^18-20^. HtrA2/Omi is evolutionarily well-conserved with respect to amino acid sequence and its three-dimensional structure^20^, suggesting that the mammalian version of HtrA/DegP could also play proteolytic roles to specifically recognize and degrade denatured proteins in mammals. In fact, the HtrA2/Omi knockout mouse and loss-of function HtrA2/Omi mutant have both demonstrated that HtrA2/Omi functions as a neuroprotective protein to prevent PD^6, 21, 22^. Accordingly, mutations in HrA2/Omi have been repeatedly found in patients suffering from PD^23-26^. However, the molecular mechanism underlying the neuroprotective role of HtrA2/Omi in PD has remained unknown until now, although it is certain that HtrA2/Omi plays an essential role in preventing PD.

Because the main function of HtrA/DegP in bacteria is to recognize misfolded or aggregated proteins to specifically degrade those proteins, we speculated that HtrA2/Omi could play an important role in removing misfolded or aggregated proteins in mammals as it does in bacteria. Because all the *in vivo* and clinical data consistently indicate that HtrA2/Omi is linked to PD progression and oligomeric α-Syn is the main misfolded protein aggregate in neurons, we investigated the potential molecular mechanism of HtrA2/Omi in terms of whether it specifically inhibits the formation of misfolded α-Syn or degrades oligomeric α-Syn to prevent PD. All of our *in vitro* and *in vivo* experiments using transgenic *Drosophila* and mice showed that HtrA2/Omi specifically recognizes and degrades oligomeric α-Syn but not monomeric α-Syn, indicating that HtrA2/Omi prevents oligomeric α-Syn-induced neurotoxicity to protect neurons from neurodegeneration by removing specifically misfolded or aggregated proteins, i.e., oligomeric α-Syn, just like it does in bacteria.

## RESULTS

### HtrA2/Omi specifically recognized and degraded oligomeric α-Syn

To confirm our speculation concerning whether the function of HtrA2/Omi in mammals is evolutionary conserved to protect neurons from oligomeric α-Syn-induced toxicity, we examined how human recombinant HtrA2/Omi (hOmi) produced in *E. coli* BL21 (DE3) pLysS-pET28a+ reacted with oligomerized α-Syn. As shown in Fig. 1a, hOmi specifically removed oligomeric α-Syn at 37 ~ 41℃ without affecting monomeric α-Syn. These data raised both possibilities that HtrA2/Omi removed oligomeric α-Syn by degradation or a chaperone action on oligomeric α-Syn to re-establish its monomeric form. To investigate these two possibilities of hOmi on oligomeric α-Syn, we specifically isolated oligomeric α-Syn from oligomerized α-Syn (Supplementary Information, Figure S1a) using a size exclusion column (Supplementary Information, Figure S1b). hOmi treatment of the purified oligomeric α-Syn resulted in complete degradation, while hOmi treatment had no effect on monomeric α-Syn (Fig. 1b). We further confirmed the oligomer-specific degradation of α-Syn by hOmi using the oligomer-specific fluorescent dye thioflavin-T (ThT). Supplementary Information Figure S2 shows that hOmi not only degraded oligomeric α-Syn specifically but also in a manner that was dose-dependent on its substrate, oligomeric α-Syn, indicating that hOmi precisely recognizes only oligomeric α-Syn. These results clearly indicated that hOmi specifically recognized and degraded oligomeric α-Syn without affecting monomeric α-Syn, a native form of α-Syn. In addition, because of the specific removal of oligomeric α-Syn by hOmi, co-treatment of oligomerized α-Syn consisting of a mixture of oligomeric and monomeric α-Syn resulted in a significant increase in cell viability in response to hOmi in a dose-dependent manner (Supplementary Information, Figure S3).

**Fig. 1.**
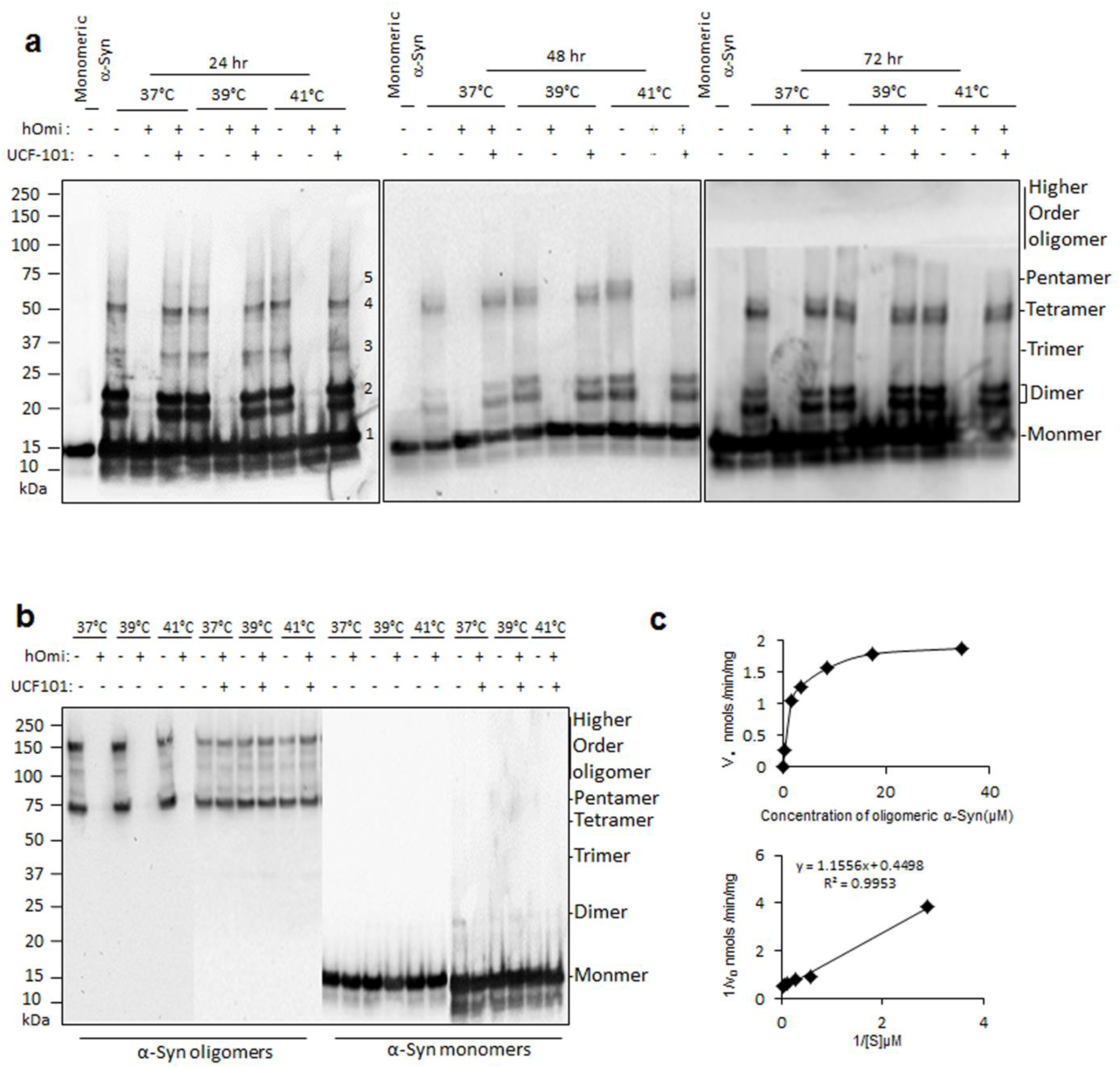
*In vitro* experiment showing that hOmi recognized and degraded specifically oligomeric α-Syn . **a** Removal of α-Syn oligomer by hOmi during the oligomerization of α-Syn at different temperatures. Treatment of UCF-101, a hOmi inhibitor, completely inhibited the oligomeric α-Syn-specific degradation activity of hOmi. **b** Complete degradation of oligomeric α-Syn without affecting monomeric α-Syn by hOmi at different temperatures but not in the presence of UCF-101. Treatment of UCF-101, a hOmi inhibitor, completely inhibited the oligomeric α-Syn-specific degradation activity of hOmi. **c** The Michaelis-Menten saturation curve (upper panel) and Lineweaver–Burk plot (lower panel) of hOmi for oligomeric α-syn. The enzyme kinetic study was conducted after labeling oligomerized α-Syn with the oligomer-specific fluorescent dye ThT.

HtrA2/Omi is an evolutionarily well-conserved serine protease, and its protease activity is inhibited by UCF-101^20^. As expected, UCF-101 completely inhibited the oligomeric α-Syn-specific protease activity of hOmi (Fig. 1a, b). These results indicated that the nucleophilic attack reaction by serine in the active site of hOmi was responsible for the oligomer-specific degradation of α-Syn. After identifying the enzymatic characteristics, we further analyzed the enzymatic kinetics of oligomeric α-Syn hydrolysis by hOmi after labeling α-Syn with ThT. The Lineweaver-Burk plot on the reactions yielded a K_m_ value of 2.569 µM and V_max_ value of 2.223 nmol/min/mg protein for oligomeric α-Syn degradation (Fig. 1c).

### Loss of HtrA2/Omi led to an accumulation of oligomeric α-Syn in mouse brain

The *in vitro* experiments examining the effects of HtrA2/Omi on oligomeric α-Syn raised questions regarding the *in vivo* role of HtrA2/Omi. Before investigating the *in vivo* functions of HtrA2/Omi, we tested whether hOmi could function as a general protease like other serine proteases, such as trypsin, or as a specific protease for particular substrates. Coincubation of hOmi with brain extracts from mnd2, HtrA2/Omi-null mutant, and wild type mice did not reveal any noticeable proteolytic degradation (Fig. 2a), which indicated that hOmi functioned as a very specific protease. Western blotting of the gel with a α-Syn-specific monoclonal antibody showed that the mnd2/mnd2 mouse accumulated a large quantity of oligomeric α-Syn, unlike the wild type littermate control (Fig. 2b). The accumulation of a large quantity of oligomeric α-Syn in mnd2 mice cast light on the pathogenic mechanism by which the mutation of HtrA2/Omi causes PD. Since oligomeric α-Syn directly causes PD^14, 17, 27^, it is reasonable that the accumulation of a large quantity of oligomeric α-Syn in mnd2 mice induces PD. hOmi treatment of the total protein extract from mnd2 mice resulted in complete degradation of the accumulated α-Syn oligomers without affecting the monomers (Fig. 2b). This result suggests that the loss of HtrA2/Omi, as in mnd2/mnd2 mice, causes PD by the loss of its ability to degrade oligomeric α-Syn.

**Fig. 2.**
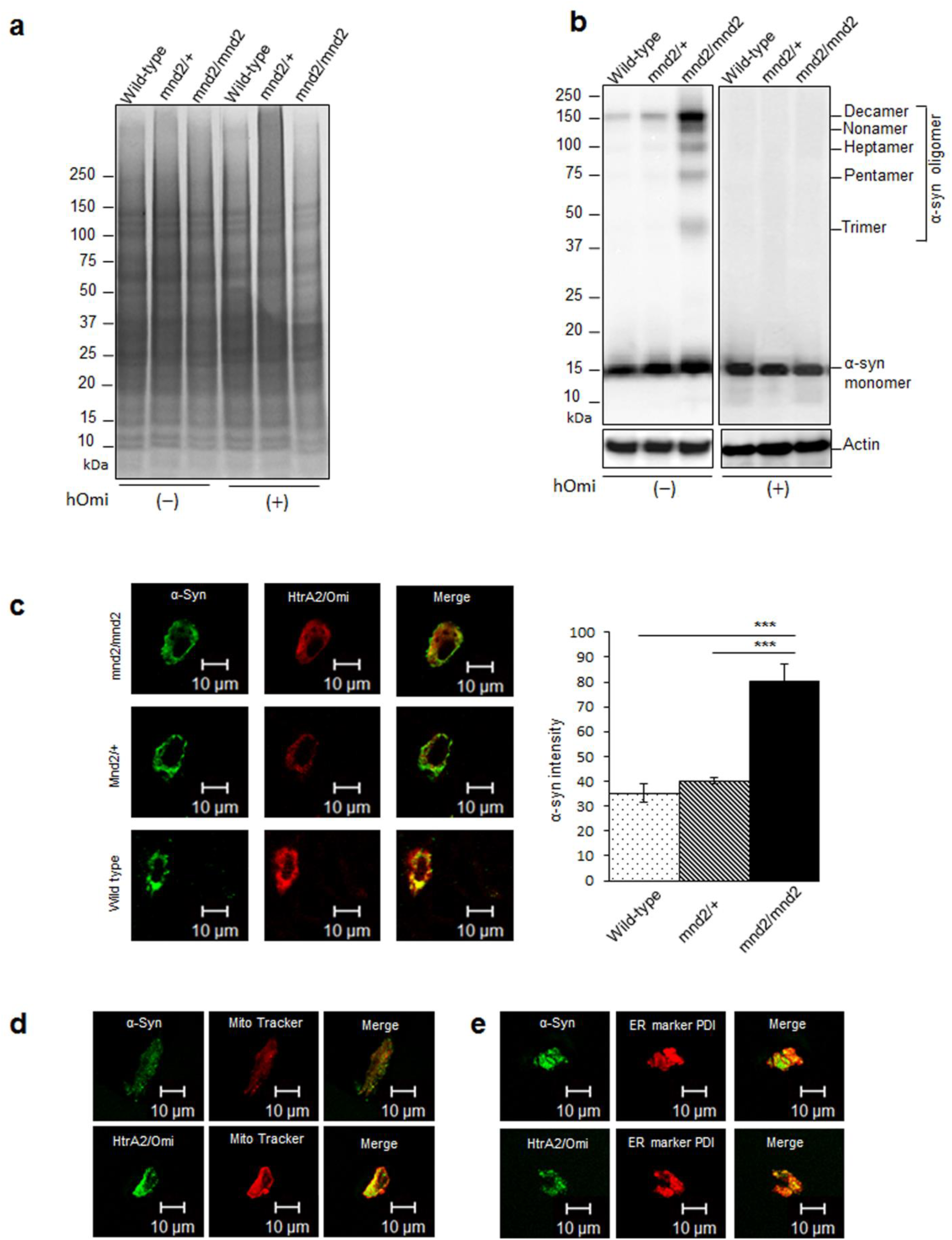
hOmi specifically recognized and degraded oligomeric α-Syn in mouse brain. **a** HtrA2/Omi treatment of total protein extracts of nigrostriatal tissues of wild-type, mnd2/+ and mnd2/mnd2 mice did not show any noticeable degradation by SDS-PAGE. **b** Western blotting with mouse anti-human α-Syn revealed a significant accumulation of oligomeric α-Syn in the nigrostriatal tissues of mnd2/nmd2 mice and complete degradation of α-Syn oligomers after hOmi treatment, without affecting monomers in the tested mice. **c** Immunohistochemical confocal microscopy of nigrostriatal tissue of the brains of wild-type, mnd2/+ and mnd2/mnd2 mice revealed the co-localization of α-Syn (green) and HtrA2/Omi (red), with significantly higher levels of α-Syn accumulation in mnd2/mnd2 mice. Representative images are shown along with the image analysis graph. **d** Immunostaining microscopy of the neurons in the substantia nigra and striatum showing the co-localization of α-Syn (green), HtrA2/Omi (green) and mitochondria (red with Mito Tracker). Scale bar, 10 µM. **e** Immunostaining microscopy of neurons in the substantia nigra and striatum showing the co-localization of α-Syn (green), HtrA2/Omi (green) and endoplasmic reticulum (red with ER marker PDI). Scale bar, 10 µM.

If the proteolytic activity of HtrA2/Omi is required to prevent oligomeric α-Syn-induced neurotoxicity, the intracellular localization of HtrA2/Omi and α-Syn should be equivalent. The immunohistochemical confocal microscopy experiments examining the substantia nigra and striatum of 4-week-old mnd2/mnd2 mice and their age-matched wild-type littermates confirmed the co-localization of HtrA2/Omi and α-Syn in mouse brain tissue (Fig. 2c), indicating that the intracellular localization of HtrA2/Omi and α-Syn is equivalent. The immunocytochemical study showed that α-Syn and HtrA2/Omi were located in mitochondria (Fig. 2d) and ER (Fig. 2e). Overall, the mnd2 mouse experiments suggested that the failure of HtrA2/Omi to remove neurotoxic oligomeric α-Syn in the ER and mitochondria led to ER stress and mitochondrial dysfunction in neurons through the accumulation of a large quantity of oligomeric α-Syn, the hallmark of PD pathogenesis. Although ER stress and mitochondrial dysfunction in neurons are the most evident pathological phenomena observed in PD^28, 29^, the mechanisms underlying ER stress and mitochondrial dysfunction in PD neurons have not been elucidated. This work clearly showed the pathological mechanism of ER stress and mitochondrial dysfunction in PD.

### Pan-neuronal expression of hOmi rescued Parkinsonism in a *Drosophila* model of Parkinson’s disease

After observing the specific degradation of oligomeric α-Syn by hOmi in mnd2 mice, we created a transgenic hOmi *Drosophila* with *w^1118^ Drosophila melanogaster* by inserting the full-length human Omi gene under the control of the UAS promoter (*UAS-hOmi*), where the heat shock 70 promoter was used as a control source of transposase (Supplementary Information, Figure S4). The transgenic hOmi *Drosophila* line was heterozygous for the dominantly marked CyO balancer chromosome carrying a dominant mutation, CyO, which causes curly wings for easy detection. From the results of the genotyping and protein expression levels of transgenic hOmi *Drosophila* flies, Tg4 (*X/Y; hOmi/Cyo;* +/+), in which *UAS-hOmi* was integrated into chromosome 2, was identified as the best hOmi transgenic line for subsequent experiments.

The hOmi *Drosophila* Tg4 was bred with a *Drosophila* model of Parkinson’s disease (α-Syn *Drosophila*) carrying the homozygous human α-Syn gene (*UAS-α-Syn*) on chromosome 3 (Supplementary Information, Figure S5). Female α-Syn *Drosophila* were mated with male hOmi *Drosophila*, and *+/hOmi; α-Syn/+* flies were selected based on the dominant phenotypes of the balancer chromosome CyO. The first filial *+/hOmi; α-Syn/+* flies were crossed to generate various genotypes. The final homozygous *x/y; hOmi/hOmi; α-Syn/α-Syn* flies were selected after genotyping the progenies from the F2 generation. The male *hOmi/hOmi; α-Syn/α-Syn* flies, 45M, were crossed with virgin female elav-GAL4 flies for pan-neuronal co-expression of hOmi and α-Syn (hOmi/α-Syn *Drosophila, x/y; +/hOmi; +/α-Syn*).

Since the Parkinsonism phenotype of α-Syn *Drosophila* is characterized by locomotor defects accompanied by reduced survivability^30, 31^, locomotor defects and survivability were tested in the hOmi, α-Syn and hOmi/α-Syn *Drosophila* lines to assess the effect of hOmi on α-Syn-induced Parkinsonism (Fig. 3). As shown in Fig. 3a, loss of climbing ability of α-Syn *Drosophila* was completely rescued by co-expression with hOmi. The performance index of locomotion was significantly lower in α-Syn *Drosophila* than hOmi/α-Syn *Drosophila* or hOmi *Drosophila* of the same age. As previously reported, the survival rate of α-Syn *Drosophila* was significantly reduced as a phenotype of *Drosophila* Parkinsonism. However, the survival rate of hOmi/α-Syn *Drosophila* was increased as much as wild type, indicating that hOmi/HtrA2 completely rescued the *Drosophila* Parkinsonism induced by α-Syn (Fig. 3b). Kaplan–Meier Survival analyses also showed that hOmi/HtrA2 completely rescued the *Drosophila* Parkinsonism (Supplementary Information, Figure S6). Overall, pan-neuronal co-expression of hOmi with α-Syn completely rescued the Parkinsonism phenotypes of α-Syn *Drosophila.* Additionally, it is interesting to note that pan-neuronal sole expression of hOmi (hOmi *Drosophila)* resulted in better performance in the locomotor reaction and increased survival rate compared with the wild type control.

**Fig. 3.**
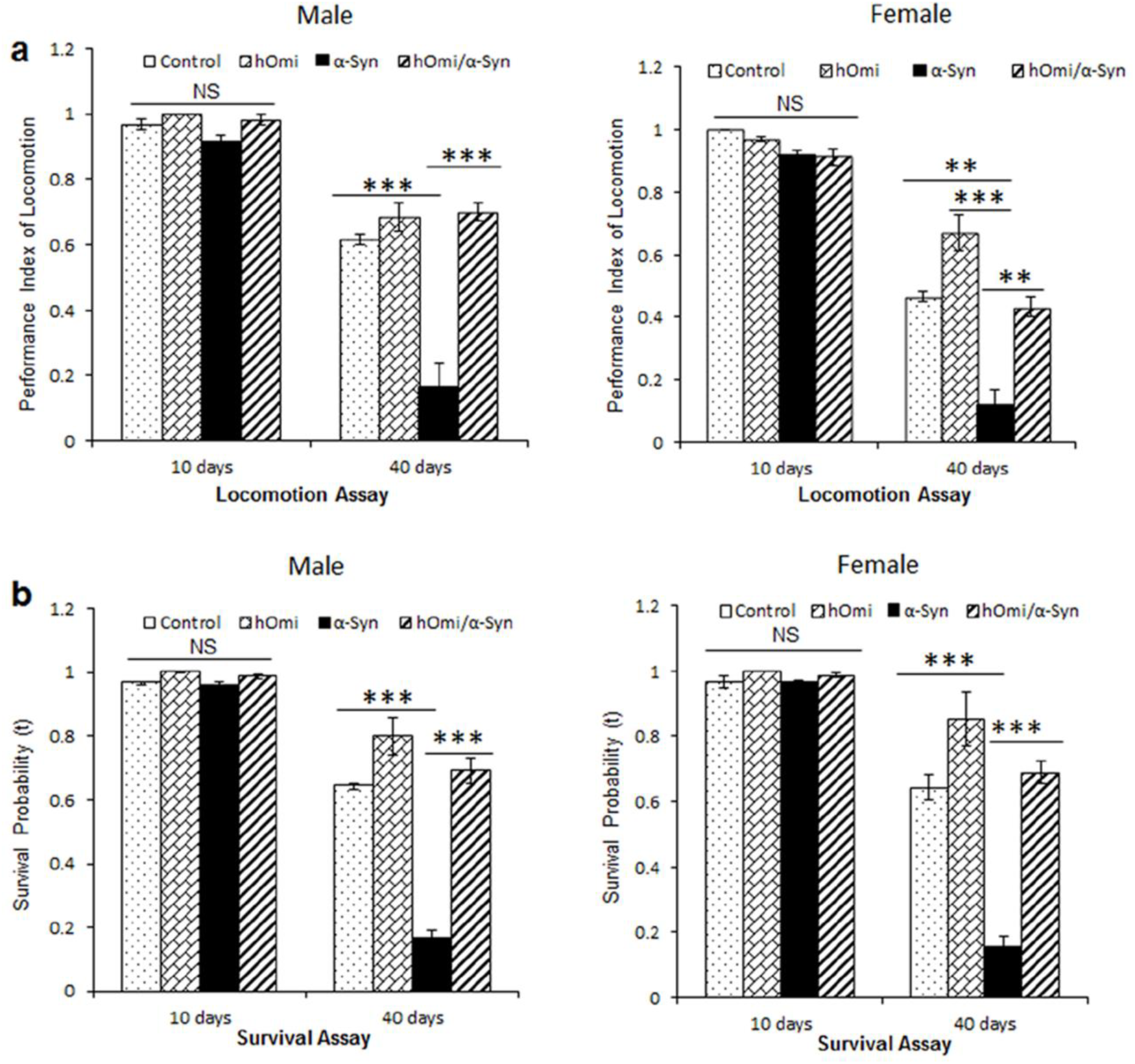
hOmi rescued Parkinsonism in a *Drosophila* Model of Parkinson’s Disease. **a** The locomotion assay in hOmi, α-Syn, or hOmi/α-Syn flies measured by climbing ability against negative geotaxis at young (10 days) and old (40 days) ages. Values are the mean ± SEM from three independent experiments. NS, not significant, ^*^*p*<0.05, *\*\*p*<0.01, ^***^*p*<0.001. **b** The survival rate of hOmi, α-Syn, or hOmi/α-Syn flies at young (10 days) and old (40 days) ages. Values are the mean ± SEM from three independent experiments. NS, not significant, ^*^*p*<0.05, *\*\*p*<0.01, *\*\*\*p*<0.001.

### Human HtrA2/Omi rescued the α-Syn-Induced neurotoxicity in a *Drosophila* model of Parkinson’s disease by oligomeric α-Syn-specific degradation

To investigate how hOmi rescues α-Syn-induced neurotoxicity in *Drosophila,* histological examinations were performed using the hOmi/α-Syn *Drosophila* line along with wild-type, hOmi and α-Syn *Drosophila* lines. Immunohistochemical confocal microscopy using an oligomer-specific monoclonal antibody, anti-α-Syn (ASy05), on brain sections showed that co-expression of hOmi and α-Syn completely eliminated the oligomeric α-Syn (Fig. 4a), which is consistent with the *in vitro* and mouse experiments (Fig. 1 and 2). Quantification of the green fluorescent intensity in the flies definitively revealed a large quantity of oligomeric α-Syn accumulation only in α-Syn *Drosophila* (Fig. 4b). However, there were no detectable α-Syn oligomers in hOmi/α-Syn *Drosophila.* We further confirmed the specific degradation of α-Syn oligomers by hOmi with total protein extracts of hOmi/α-Syn fly brains (Fig. 4c). Anti-α-Syn antibody detected both oligomeric and monomeric α-Syn in α-Syn *Drosophila*. However, only monomeric α-Syn was detected by western blotting of hOmi/α-Syn *Drosophila*. This *in vivo* result clearly confirmed that HtrA2/Omi specifically recognized and degraded oligomeric α-Syn without affecting monomeric α-Syn. Considering that oligomeric α-Syn has strong neurotoxicity to function as an etiological agent for PD while monomeric α-Syn lacks neurotoxicity, rather playing an essential role in maintaining a supply of synaptic vesicles in presynaptic terminals^32, 33^, this result shed light on how hOmi provides a neuroprotective function in PD.

**Fig. 4.**
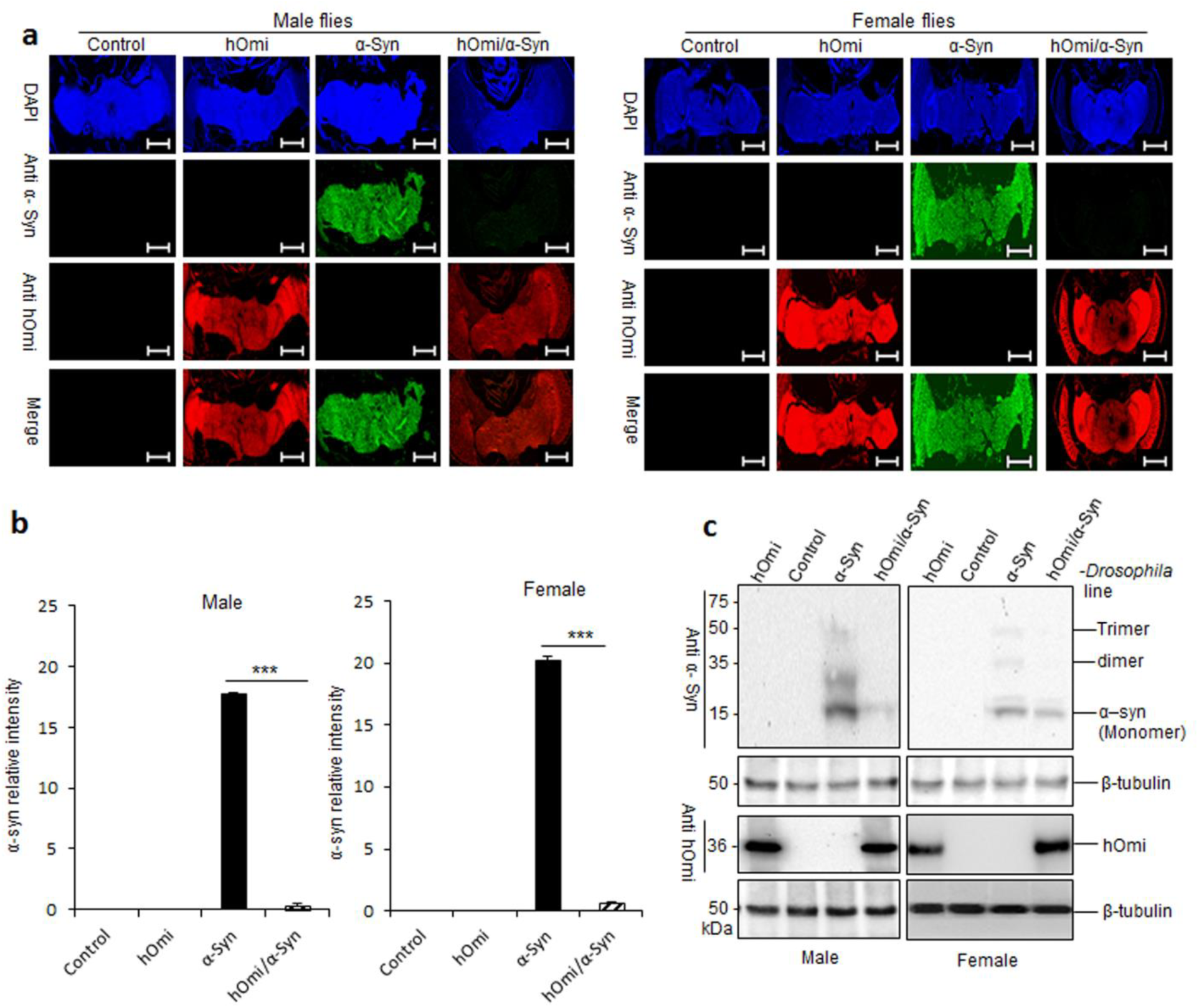
hOmi prevented the accumulation of oligomeric α-Syn in a *Drosophila* Model of Parkinson’s Disease. **a** Immunohistochemical confocal microscopy of the brains of control flies and transgenic flies expressing hOmi (indicated in red with anti-hOmi), α-Syn (indicated in green with anti-α-Syn) and hOmi/α-Syn using 40-day-old male (left panel) and female flies (right panel). The α-Syn was stained with an oligomeric α-Syn-specific monoclonal antibody, anti-α-Syn (ASy05). Scale bar, 100 µM. **b** Relative intensity of oligomeric α-Syn immunofluorescence in the images in a. Values are the mean ± SEM from three independent experiments. ^*^*p*<0.05, ^**^*p*<0.01, *\*\*\*p*<0.001. **c** Western blot analysis of α-Syn and hOmi expression in Drosophila brains. The blots of the brain homogenates from control flies and transgenic flies expressing hOmi, α-Syn or hOmi/α-Syn using 40-day-old male (left panel) and female flies (right panel) were probed with anti-α-Syn or anti-hOmi.

We further investigated *Drosophila* brains after immunostaining with anti-α-Syn and anti-HtrA2 antibodies. An age-dependent accumulation of α-Syn clearly caused the accumulation of Lewy bodies in α-Syn *Drosophila*, whereas co-expression of hOmi with α-Syn in hOmi/α-Syn *Drosophila* completely prevented the accumulation of Lewy bodies, and the overall integrity of the brain tissue was the same as the normal control (Fig. 5a, b). H&E staining of the brain slices of the flies further confirmed a clear neurodegeneration in the α-Syn *Drosophila* (Fig. 6a, b). The integrity of the brain tissue was observed in both young and aged hOmi/α-Syn *Drosophila* as in the case of control flies and hOmi *Drosophila*, but not in aged α-Syn *Drosophila* due to neurodegeneration. The brains of 40-day-old α-Syn *Drosophila* showed clear neuronal loss with astrocytosis and the appearance of Lewy bodies both in male (Fig. 6a) and female flies (Fig. 6b). In accordance with the previous results concerning the function of hOmi in oligomeric α-Syn-specific degradation, this result again confirmed that hOmi rescued the α-Syn-induced neurotoxicity in α-Syn *Drosophila*.

**Fig. 5.**
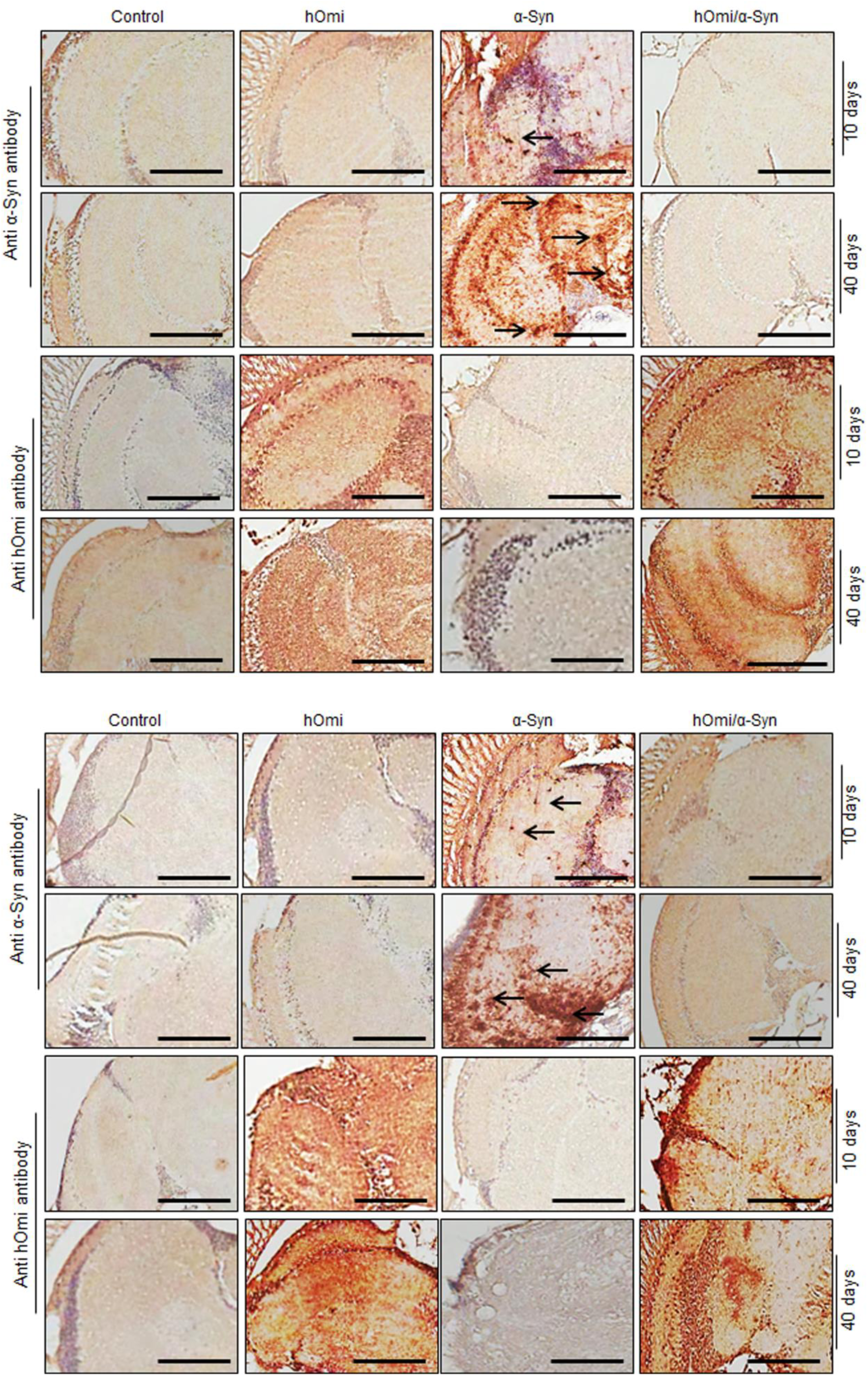
hOmi prevented the formation of Lewy bodies and maintained brain integrity in a *Drosophila* Model of Parkinson’s Disease. **a** Immunohistochemical staining of the midbrains of male control flies and transgenic flies expressing hOmi, α-Syn and hOmi/α-Syn with either anti-α-Syn or anti-hOmi antibody, showing Lewy bodies (indicated as arrows) in 10 and 40-day-old male flies. The α-Syn was stained with an oligomeric α-Syn-specific monoclonal antibody, anti-α-Syn (ASy05). Scale bar, 50 µM. **b** Immunohistochemical staining of the midbrains of female control flies and transgenic flies expressing hOmi, α-Syn and hOmi/α-Syn with either anti-α-Syn or anti-hOmi antibody, showing Lewy bodies (indicated as arrows) in 10 and 40-day-old female mice. The α-Syn was stained with an oligomeric α-Syn-specific monoclonal antibody, anti-α-Syn (ASy05). Scale bar, 50 µM.

**Fig. 6.**
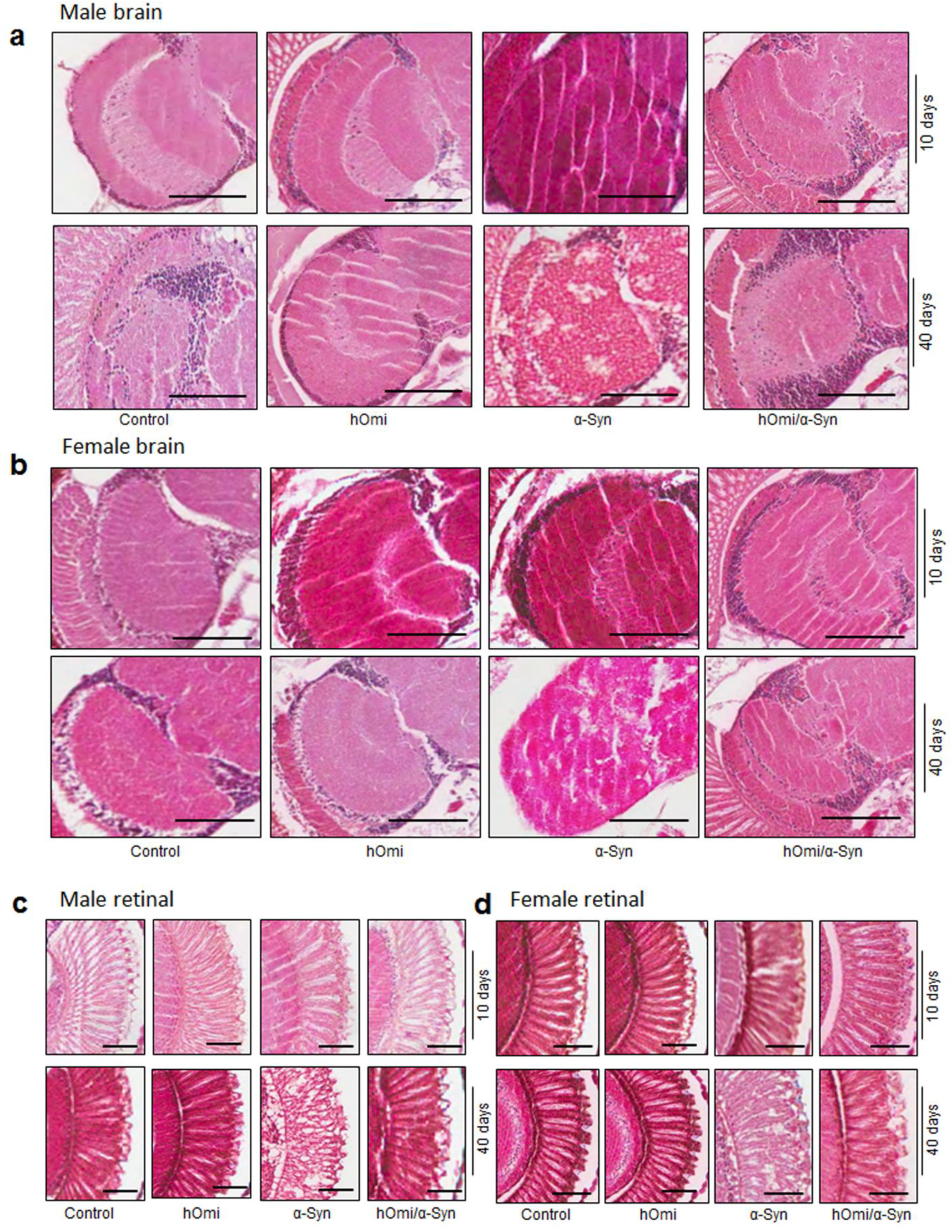
Histological examination confirming that hOmi prevented α-Syn-Induced neurodegeneration in a Drosophila Model of Parkinson’s Disease. **a** H&E staining of the cortex and neuropil region of male control flies and transgenic flies expressing hOmi, α-Syn, or hOmi/α-Syn, aged 10 or 40 days. Scale bar, 50 µM. **b** H&E staining of the cortex and neuropil region of female control flies and transgenic flies expressing hOmi, α-Syn, or hOmi/α-Syn, aged 10 or 40 days. Scale bar, 50 µM. **c** H&E staining of retinal sections of male control flies and transgenic flies expressing hOmi, α-Syn, or hOmi/α-Syn, aged 10 days or 40 days. Scale bar, 50 µM. **d** H&E staining of retinal sections of female control flies and transgenic flies expressing hOmi, α-Syn, or hOmi/α-Syn, aged 10 days or 40 days. Scale bar, 50 µM.

### Human HtrA2/Omi counteracted the α-Syn-induced developmental defect in *Drosophila* eye

It has been previously observed that the expression of α-Syn in the developing eye causes retinal degeneration in *Drosophila*^34^. Since α-Syn-induced retinal degeneration well-represented α-Syn-induced neurotoxicity, we crossed the GMR-GAL4 driver line with the α-Syn, hOmi and hOmi/α-Syn *Drosophila* lines to drive the expression of transgenes in the ommatidial unit and selected the transgene-expressed flies based on the dominant phenotype of the balancer chromosome CyO of GMR-GAL4. The eye-specific expression of α-Syn clearly revealed degenerated retina (Fig. 6c, d). As the α-Syn *Drosophila* aged from 10 to 40 days, substantial vacuolar changes became evident, which indicated that α-Syn acted as an etiological agent for retinal degeneration. In contrast, co-expression of hOmi with α-Syn in hOmi/α-Syn *Drosophila* did not show any retinal degeneration in either male (Fig. 6c) or female flies (Fig. 6d).

Additionally, the expression of α-Syn led to developmental defects of the eyes, showing a loss of general retinal tissue integrity and roughness of the eye (Fig. 7). Serious eye defects were observed in both male and female α-Syn *Drosophila*, and the eye defects became more serious as the flies aged (bottom panel of Fig. 7a, b). In contrast, hOmi/α-Syn *Drosophila* did not show any eye defects and the eye phenotype was equivalent to the normal control (GMR-GAL4) and hOmi *Drosophila* (Fig. 7a, b). Scanning electron microscopy also revealed serious defects of the α-Syn *Drosophila* eye and the normal undamaged eye of hOmi/α-Syn *Drosophila* (Fig. 7c and 7d). Ommatidial disarray was significantly increased in α-Syn *Drosophila* compared with hOmi/α-Syn *Drosophila,* hOmi *Drosophila* or the normal control (GMR-GAL4), and the difference became more evident as the flies increased in age (bottom panel of Fig. 7c, d). Furthermore, the bristle of the eye of α-Syn *Drosophila* was predominant and became seriously lost as the flies aged. However, this phenomenon was not observed in hOmi/α-Syn *Drosophila,* hOmi *Drosophila* or the normal control (GMR-GAL4) (Fig. 7e, f). This result clearly demonstrated the neuroprotective role of hOmi in α-Syn-induced neurodegeneration.

**Fig. 7.**
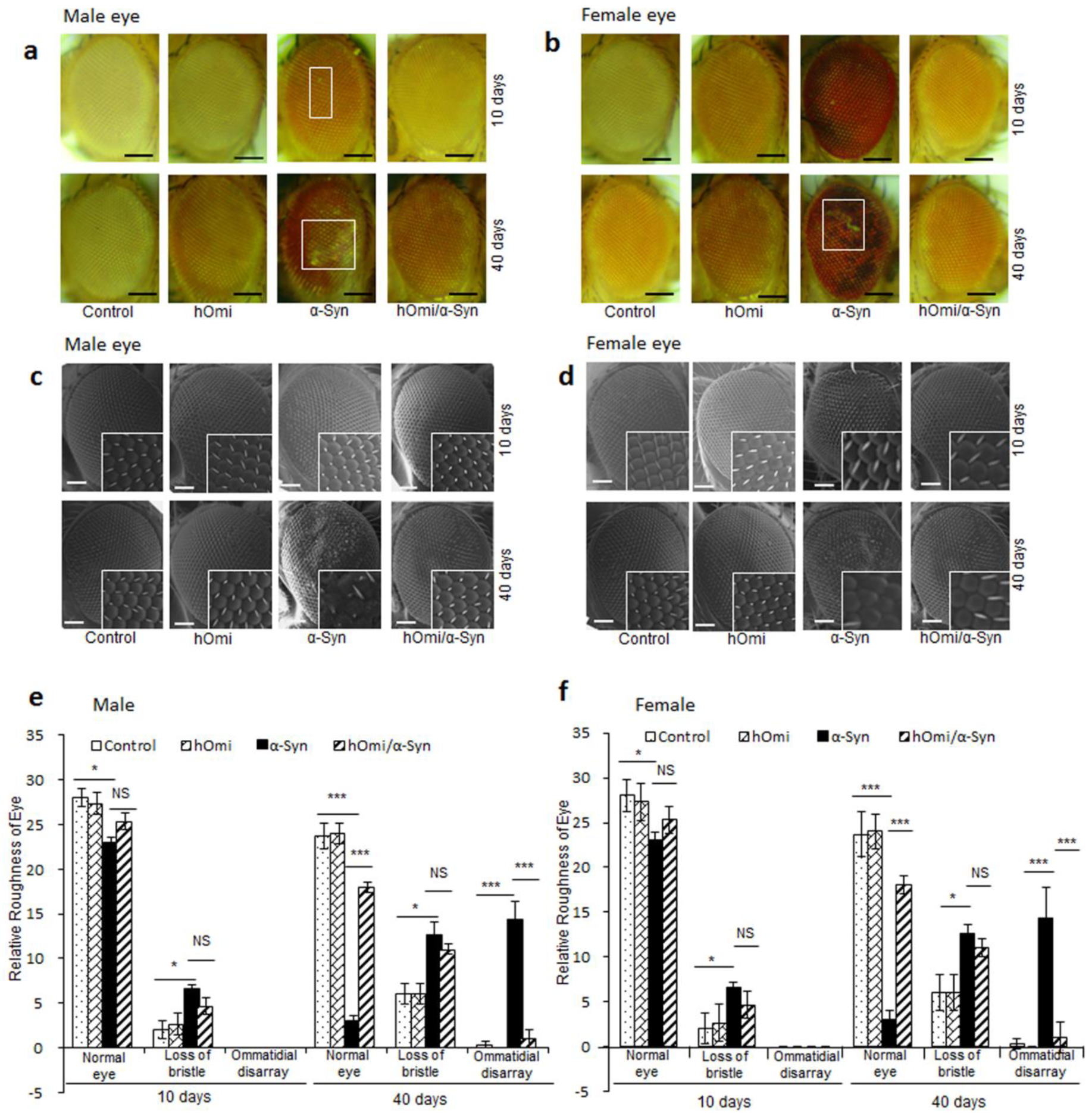
hOmi counteracted the α-Syn-induced developmental defects in *Drosophila* Eye. **a, b** Light microscopic images of the eyes of control and transgenic flies expressing hOmi, α-Syn, or hOmi/α-Syn, aged 10 days or 40 days in male a and female flies b. Scale bar, 100 µM. **c, d** Scanning electron microscopy images of eyes of control and transgenic flies expressing hOmi, α-Syn, or hOmi/α-Syn, aged 10 days or 40 days in male c and female flies d. Scale bar, 50 µM. The 4 × magnifications are presented in the square box. **e, f** Roughness counting of eye phenotypes based on normal phenotypes, loss of bristles and ommatidial disarray of control and transgenic flies expressing hOmi, α-Syn, or hOmi/α-Syn, aged 10 days or 40 days in male e and female flies f. Values are the mean ± SEM from three independent experiments. NS, not significant, ^*^*p*<0.05, *\*\*p*<0.01, ^***^*p*<0.001.

## DISCUSSION

### Dichotomous reports of HtrA2/Omi

HtrA2/Omi is a mitochondrial protein with high homology to a bacterial heat shock protein^18, 19^. Therefore, it was speculated that HtrA2/Omi would function similarly to the bacterial protein to protect cells from stress-induced toxicity caused by misfolded proteins. Despite its protective role in bacteria, *in vitro* studies have shown that HtrA2/Omi acts as a pro-apoptotic protein^35^. HtrA2/Omi is released into the cytosol from mitochondria during apoptosis and degrades inhibitor of apoptosis proteins (IAPs) such as XIAP and CIAP1/2^35-37^. The degradation of these IAPs by HtrA2/Omi activates both caspase-dependent and -independent apoptotic pathways^36, 37^.

Considering the pro-apoptotic characteristics of HtrA2/Omi, it would be natural to think that HtrA2/Omi could participate in a disease-escalating process rather than a disease-protecting process. However, *in vivo* animal experiments from both mice and insects have shown that HtrA2/Omi does not play a pro-apoptotic role, in contrast to the *in vitro* findings^20, 38^. Rather, HtrA2/Omi is not only dispensable for apoptosis but also allows brains to be maintained a healthy state. This work showed the same basic trend as the previous results of *in vivo* HtrA2/Omi expression experiments. As shown in Fig. 3~6, expression of HtrA2/Omi in the *Drosophila* brain not only maintained the health and integrity of the brain but also increased life span. Thus, hOmi *Drosophila* showed demonstrated that the functions of HtrA2/Omi are essential for maintaining the health of the brain.

Although apoptosis is the end of life to cells, some apoptosis is required to maintain the healthy state of multicellular organisms. In this context, the paradoxical results obtained for HtrA2/Omi *in vitro* and *in vivo* provide an abstruse example of the life process of multicellular organisms.

### Significance of oligomeric α-Syn-Specific degradation by HtrA2/Omi in the etiology of PD

In accordance with pan-neuronal expression experiments of HtrA2/Omi *in vivo*, the knock-out mouse of HtrA2/Omi and the natural HtrA2/Omi mutant mouse demonstrated that HtrA2/Omi functions as a neuroprotective protein to prevent PD^21, 39^. The presence of mutations/polymorphisms in HtrA2/Omi in sporadic PD patients further solidified the link of HtrA2/Omi to PD^22^. Due to the functional loss of HtrA2/Omi *in vivo* and clinical observations showing an association of HtrA2/Omi with PD, HtrA2/Omi was named PARK13 to represent a PD gene.

Because mammalian Omi/HtrA2 binds to PINK1 and the phosphorylation of Omi/HtrA2 is dependent on PINK1^40^, it has been suggested that HtrA2/Omi functions downstream of the PINK1/Parkin pathway. However, extensive loss-of-function-based genetic interaction studies using *Drosophila* have failed to show an association of Omi/HtrA2 either upstream or downstream of PINK1^31, 41^. Gene interaction studies using mice have also shown that overexpression of Parkin does not rescue neurodegeneration in the Omi/HtrA2 mutant^42^. These studies clearly indicate that HtrA2/Omi does not function in the PINK1/Parkin pathway. Although HtrA2/Omi certainly plays a neuroprotective role to prevent PD, the functional mechanism of HtrA2/Omi has remained mysterious until now. The gene interaction studies in this work clearly showed that HtrA2/Omi specifically degraded only the neurotoxic form of α-Syn, oligomeric α-Syn, without affecting nontoxic normal α-Syn (monomeric α-Syn).

Complete rescue of oligomeric α-Syn-induced toxicity *in vivo* sheds light on the etiology of PD. Since PINK1 certainly phosphorylates HtrA2/Omi, there is a possibility that PINK1 may be involved in the regulation of HtrA2/Omi, although HtrA2/Omi does not function in the PINK1/Parkin pathway. Thus far, thirteen genes that cause PD have been identified. The etiopathogenic mechanism of PD involving these genes can be grouped into two pathways: disruption of PINK1-associated phosphorylation in mitochondria and neurotoxic protein aggregation associated with α-Syn. Because both *in vitro* and genetic studies have suggested that HtrA2/Omi functions downstream of PINK1^26^, our results could provide a key piece of the PD puzzle that links these two pathways at the molecular level.

### The evolutionarily conserved function of HtrA2/Omi

The results of this study provide new findings about the neuroprotective role of HtrA2/Omi based on its ability to detoxify neurotoxic oligomeric α-Syn in PD. α-Syn is degraded by autophagy and the proteasome^43^; however, these degradation pathways also degrade non-amyloidogenic monomeric α-Syn, indicating that they are not related to the etiopathogenesis of PD. The oligomeric form of α-Syn is known to be resistant to all proteases, including proteinase K^44^, and the clearance mechanism for amyloidogenic α-Syn has remained unknown. Our results show that oligomeric α-Syn is specifically degraded in neurons by HtrA2/Omi to prevent PD. HtrA2/Omi not only plays a critical role in the prevention of PD, but our results are also in good agreement with the previous observation that HtrA2/Omi functions as a chaperone to detoxify oligomeric Aβ into monomeric Aβ^20^. Thus, HtrA2/Omi might be a key protein that relieves the stress caused by various amyloidogenic neuronal proteins such as oligomeric α-Syn and oligomeric Aβ.

It is well-known that the functions of most proteins are evolutionarily conserved. Bacteria have HtrA that specifically degrades misfolded proteins through its protease activity^45^. The bacterial homolog of HtrA, HtrA2/Omi, functions as a protease to specifically degrade a type of misfolded protein, *i.e.,* oligomeric α-Syn. Considering that the original function of HtrA was to degrade misfolded protein through its protease activity, it is very interesting to note that the function of the mammalian version of HtrA, HtrA2/Omi, is to remove oligomeric α-Syn through its protease activity. Because oligomeric α-Syn is the misfolded version of native α-Syn, the original function of HtrA seems to be perfectly conserved in mammals.

## MATERIALS AND METHODS

### Reagents and antibodies

The reagents used in all experiments and antibodies used for western blot or immunohistochemical analysis are listed in the Supplementary Information Table S1.

### α-Syn *Drosophila melanogaster* and driver lines for transgene expression

The α-Syn Transgenic fly UAS-α-Syn was purchased from the Bloomington *Drosophila* stock center (FBst0008146), and the transgenic hOmi *Drosophila* was created in this work. The driver line elav-GAL4 of *Drosophila melanogaster* (FBst0000458) was used for pan-neuronal expression of transgenes, and the driver line GMR-GAL4 of *Drosophila melanogaster* (FBti0002994) was used to express transgenes in the eye.

### Recombinant hOmi protein expression and purification

The plasmid construct encoding hOmi (134-458) was generated using the pET28a^+^ vector (Novagen Merck Millipore, cat# 69864-3CN). *E. coli* BL21 (DE3) pLysS (Stratagene California, cat# CMC0018) was transfected with the construct by electroporation. A single colony of *E. coli* BL21 (DE3) pLysS-pET28a^+^-HtrA2/Omi was grown at 37℃/250 rpm in 1 L LB medium containing 50 µg/mL until the OD reached ~ 0.8. One millimole of IPTG was added to the culture, followed by further culture for 5 hr at 20℃/250 rpm to induce protein expression. After induction of heterologous protein, the bacterial cells were centrifuged at 6,000 × *g* for 15 min and stored at either -20℃ or at -80℃ until further use. The harvested bacterial pellet from the 1-L culture was washed once with PBS and resuspended in 20 mL of native cell lysis buffer A (50 mM Tris-HCl, pH 8.0, 150 mM NaCl, 2.5 mM EDTA, 0.5% Triton X-100, 4 mM MgCl_2_, 50 µg/mL DNase I, 0.5 mg/mL lysozyme and protease inhibitor cocktail (Roche Applied Science, cat# 11836153001)). After incubation at room temperature for 30 min, the lysate was subjected to sonication 10 times for 10 sec with a burst speed of 6 at high intensity with a 1 min cooling period on ice using the ultrasonic homogenizer (Bandelin Sonopuls HD 2070, Berlin, Germany). Following sonication, 20 mL of denaturing buffer B (50 mM Tris-HCl, pH 8.0, 10 M urea, 500 mM NaCl and 20 mM imidazole) was added to the above bacterial cell lysate, followed by sonication 10 times for 10 sec with a burst speed of 6 at high intensity with a 1 min cooling period on ice. Thereafter, the bacterial cell lysate was centrifuged at 20,000 × *g* for 1 hr at 4℃, and the supernatant was allowed to solubilize at room temperature overnight with constant stirring on a magnetic stirrer. The solubilized protein solution was centrifuged at 20,000 × *g* for 1 hr to remove the insoluble materials, followed by an incubation at 4℃ for 1 hr with gentle agitation to bind to a 5-mL Ni-NTA agarose affinity column (Invitrogen, cat# R90101) pre-equilibrated with 30 mL of protein denaturing buffer C (50 mM Tris-HCl, pH 8.0, 5 M urea, 300 mM NaCl) containing 10 mM imidazole. After binding, the column was washed sequentially with 30 mL and 75 mL of buffer C containing 10 mM and 50 mM imidazole, respectively. The denatured protein was then eluted with three column volumes of buffer C containing 500 mM imidazole. The Ni-NTA fractions were re-purified on a PD-10 column (Amersham Pharmacia Biotech, Bellefonte, USA) equilibrated with buffer C, followed by elution with the same buffer according to the manufacturer’s instructions. After determining the protein concentration using the Bradford assay kit (Thermo Fisher Scientific, cat# 23200), the protein was reduced by DTT to a final concentration of 10 mM at 37℃ for 1 hr. Thereafter, the protein was refolded in optimized protein refolding buffer D (50 mM Tri-HCl, pH 8.5, 500 mM NaCl, 5 mM EDTA, 5 mM GSH, 0.5 mM GSSG, 500 mM arginine and 15% glycerol) using a rapid-dilution method by maintaining the final protein concentration at 40-50 µg/mL with constant stirring overnight at 4℃. Soluble refolded protein was concentrated at 4℃ on an Amicon stirred cell ultrafiltration unit using an YM-10 membrane (Millipore Sigma Aldrich, cat# Z648078). Further purification of the refolded protein was performed on a PD-10 column equilibrated with buffer E (20 mM HEPES, pH 7.5, 100 mM NaCl). The protein was eluted using the same buffer, and the purified protein concentration was stored at -80℃ in 20% glycerol.

### Generation of *Drosophila melanogaster* expressing hOmi pan-neuronally

The cDNA encoding human *Omi (hOmi)* was amplified by PCR from cDNA clones (Invitrogen Thermo Fisher, Cat# 11262011). The amplified PCR product was digested by flanking restriction enzyme sites and subcloned into the same restriction sites in the GAL4-responsive pUAST expression vector, a gift from J.C. Moon^5^. The vector construct was co-injected into *w^1118^ Drosophila melanogaster* embryos with a plasmid bearing P element transposase under the control of the heat shock 70 (hs-π) promoter as a source of transposase following standard microinjection methods (BestGene Inc., Chino Hills, CA). Balanced activator lines were generated using standard genetic techniques at BestGene Inc. The generation of transgenic flies was confirmed by PCR using primers to detect the *hOmi* gene and western blot analysis using an anti-HtrA2/Omi antibody (Supplementary Information, Fig. S4). The transgenic hOmi *Drosophila* line was heterozygous for the dominantly marked CyO balancer chromosome carrying a dominant mutation, CyO, which causes curly wings.

### Generation of *Drosophila melanogaster* co-expressing hOmi and α-Syn pan-neuronally

Female *Drosophila* model of Parkinson’s disease (α-Syn *Drosophila*, FBst0008146) was mated with male hOmi *Drosophila* which maintains hOmi with CyO balancer chromosome. In the F1 generation, the flies were sorted based on the dominant phenotypes of the balancer chromosome CyO and selected *+/hOmi; α-Syn/+* flies. The first filial *+/hOmi; α-Syn/+* flies were crossed with each other to generate various genotypes. By genotyping the progenies born from a male fly and a female fly of the F2 generation, homozygous x/y; *hOmi/hOmi; α-Syn/α-Syn Drosophila melanogaster* were identified. The identified homozygous x/y; *hOmi/hOmi; α-Syn/α-Syn Drosophila melanogaster* were maintained by crossing them with each other. The homozygous *hOmi/hOmi; α-Syn/α-Syn* male *Drosophila melanogaster* was crossed with 3~4 female elav-Gal4 virgin *Drosophila melanogaster* to produce *Drosophila melanogaster* with pan-neuronal co-expression of hOmi and α-Syn. After 48 hrs of breeding, the flies were transferred to a fresh tube. The presence of the *hOmi* and *α-Syn* genes was observed in the progeny. The genotypes of the *Drosophila* lines were detected by PCR using EF Taq Polymerase (Solgent, cat# SEF 16-R250) following the manufacturer’s suggestion, in which 5 µL EF Taq buffer, 1 µL NTP, 0.5 µL forward primer (10 pmol/µL), and 0.5 µL reverse primer (10 pmol/ µL) together with 50 ng genomic DNA in ddH_2_O water to 50 µL made up the PCR mix. *Drosophila* genomic DNA was isolated using DNAzol (Invitrogen, cat# 10503-027). After the initial denaturation at 95℃ for 5 min, PCR was carried out by denaturation at 95℃ for 30 sec, annealing at 57℃ for genotyping *α-Syn*, 65℃ for genotyping *hOmi* for 45 seconds and extension at 72℃ for 1 min. After completion of 34 cycles, a final extension of 10 minutes was applied. The PCR products were confirmed by agarose gel electrophoresis using loading star (Dyne Bio, cat# A750) and an 100-bp DNA ladder (Dyne Bio, cat# A751). α-Syn F primer: 5’-TGT AGG CTC CAA AAC CAA GG-3’; R primer: 5’-GCT CCC TCC ACT GTC TTC TG-3’ and hOmi F primer: 5’- GTC GCC GGA TCC ATG CGC TAC ATT-3’ R primer: 5’-GAG CTC TCG AGT CAT TCT GTG ACC-3’.

### Mouse Model

This mouse study was carried out in strict accordance with the recommendations of the Guide for the Ethics Committee of Chonbuk National University Laboratory Animal Center. The protocol was approved by the Ethics Committee of Chonbuk National University Laboratory Animal Center (Permit Number: CBU 2012-0040). All efforts were made to minimize suffering. The C57BL/6J-mnd2 mice (RRID: IMSR_JAX:004608) carrying a mutation at S276C in HtrA2/Omi, a Parkinson model mouse, were obtained from the Jackson Laboratory (Bar Harbor, Maine). Homozygous (mnd2/mnd2), heterozygous (mnd2/+) and wild-type mice were obtained by crossing mnd2 heterozygous (mnd2/+) mice. The genotypes of the mice were identified by PCR.

### SDS-PAGE and Western blotting

The protein samples were mixed with NuPAGE 4 × LDS sample buffer (Invitrogen, cat# NP007), heated at 95℃ for 5 min and run on a 4-12% Bis-Tris gradient gel (Invitrogen, cat# NP#00322). The proteins were visualized by staining with Coomassie blue. For western blotting, the SDA-PAGE gels were transferred to PVDF membranes (Thermo Fisher Scientific, cat# 88018). After blotting, the PVDF membranes blocked with 5% non-fat dry milk (Bio-Rad Laboratories, cat# 170-6404) in TBST (TBS with 0.05% Tween and 0.1% Triton X-100) for 2 hrs at room temperature. The membranes were washed with TBST and incubated overnight at 4℃ after adding mouse anti-α-Syn (Abcam, cat# Ab1903) at a 1:2000 dilution or mouse anti-HtrA2/Omi antibody (BD Biosciences, cat# ABIN121159) at a 1:2000 dilution. The membranes hybridized to the primary antibody were washed with TBST, followed by incubation for 2 hrs at room temperature after addition of a horseradish peroxidase-conjugated goat anti-mouse IgG (H+L) antibody (1:3000) (Promega Corporation, cat# W4021). After washing with TBST, the chemiluminescent substrate, Immun-Star^TM^ Western CTM Kit (Bio-Rad Laboratories, cat# 170-5061), was added to the membranes, and images were captured with an XRS camera equipped with a Bio-Rad Quantity One imaging system. The stock solutions of primary (anti-α-Syn and anti-HtrA2/Omi) and secondary antibodies (goat anti-mouse) were diluted with antibody dilution buffer (TBST-Triton X-100 with 0.5% BSA).

### *In vitro* enzymatic assay of HrA2/Omi

Commercially purchased human recombinant α-Syn protein (r-Peptide, cat# S-1001-1) was used in this work. For the preparation of oligomeric α-Syn, human recombinant monomeric α-Syn was diluted in NaP buffer, pH 7.4, to a final concentration of 10 µg/mL and incubated at 37℃. Ten microliters of recombinant hOmi (10 µg/mL in 50 mM sodium phosphate buffer, pH 7.4) produced in *E. coli* BL21 (DE3) pLysS-pET28a+ was incubated for 30 min at room temperature prior to addition of the same volume of human recombinant α-Syn protein (10 µg/mL). The reaction mixture was incubated at 37℃, 39℃ and 41℃ each. UCF-101 (Merck Millipore, cat# 496150), and a hOmi specific inhibitor was added to the reaction mixtures if necessary. See the Supplementary information for detailed methods. For further degradation analysis, oligomers and monomers were purified from *in vitro* oligomerized α-Syn protein using a PD-10 column using Sephadex® G-25M resin (Sigma Aldrich, cat# G25150) according to the manufacturer’s suggestions. Briefly, the dry powder of Sephadex was swollen in water overnight prior to use at 4℃. A 1-mL micropipette tip was used for bedding, and the opening of the pipette was plugged with glass wool (Sigma Aldrich, cat# 20411). The column was gently poured down the side of the pipette. The protein was eluted with sodium phosphate buffer. The purity of the purified monomeric and oligomeric α-Syn was checked by western blotting using a mouse anti-α-Syn antibody (Abcam, cat# Ab1903). Thereafter, following the same procedure, an equal volume of purified α-Syn oligomer or monomer, 10 µg/mL, was incubated with the recombinant hOmi. The reaction mixtures were analyzed by western blotting using a mouse anti-α-Syn antibody. See the Supplementary information for SDS-PAGE & Western blot detailed methods.

### Enzymatic kinetics of HtrA2/Omi using ThT

To assess the enzymatic kinetics of hOmi, 5 µM of oligomeric α-Syn was incubated with different concentration of hOmi (0 nM, 10 nM, 50 nM, and 100 nM) in 150 µL of 10 µM ThT (Sigma Aldrich, cat# 2390-54-7) solution at room temperature using sealed 96-well plates. The ThT fluorescence intensity of each sample was measured at 485~540 nm every 5 min using an iMark™ microplate reader (Bio-Rad, Hercules, CA). The ThT fluorescence was also measured to obtain the Lineweaver-Burk plot. The plot of the reactions was used to calculate K_m_ and V_max_ values for hOmi.

### Cell viability assay

The cell viability was evaluated using the Cell Counting Kit-8 (Sigma Aldrich, cat # 96992). Primary mouse neurons were harvested, and 2 × 10^5^ neurons were plated in 24-well polystyrene plates. The plates were incubated at 37℃ for 24 hrs for neuronal attachment. After 24 hrs, primary mouse neurons were treated with α-synuclein. The plates were then incubated at 37℃ for 3 hrs, 24 hrs and 48 hrs. After incubation, 10 μL of the reaction solution was added to each well and incubated for 4 hr at 37°C according to the manufacturer’s instructions. The absorbance of each well was then measured at 450 nm using a microplate reader (Bio-Rad Laboratories, Hercules, California). All experiments were repeated at least three times.

### Histological examinations of the brains of mice and *Drosophila*

Brains of four-week-old mnd2/mnd2 mice and age-matched control mice were removed and fixed by transcardial perfusion with ice-cold PBS followed by 4% paraformaldehyde in PBS. Subsequently, those were post-fixed in the same fixative for 4 hrs at 4℃ and incubated overnight at 4℃ in 30% sucrose in PBS. For the Drosophila experiment, fly heads were fixed in 10% neutral-buffered formalin. The fixed mouse brains and fly heads were embedded in paraffin and sliced into 4-μM-thick sections on a freezing microtome machine. The sections were deparaffinized with xylene, rehydrated with a descending series of diluted ethanol and water, and permeabilized with 1% Triton X-100 in TBS for 30 min at RT. Antigens in the sections were retrieved with 10 mM sodium citrate buffer (pH 6.0) for 20 min at 85℃ and subsequently blocked with 10% normal goat serum (Sigma Aldrich, cat# G9023) in TBS containing 1% BSA and 0.025% Triton X-100 at RT for 10 hrs. To observe the co-localization of α-Syn and hOmi in mouse brain, the sections were double-stained at 4℃ with a mouse anti-α-Syn antibody (Abcam, cat# Ab1903) and a rabbit anti-HtrA2/Omi antibody (1:200) (Abcam, cat# Ab64111). For the *Drosophila* brain, however, the sections were double-stained at 4℃ with mouse anti-α-Syn antibody ASy05 (1:400) (Agricera antibodies, cat# AS132718) and a rabbit anti-HtrA2/Omi antibody (1:400) (Abcam, cat# Ab64111). The sections were sequentially washed 3 times for 10 min with TBST (TBS and 0.20% Tween-20), 2 times for 5 min with TBS-Triton X-100 (TBS and 0.025% Triton X-100) and one time for 10 min with TBST. To suppress endogenous peroxidase activity, the sections were incubated with 3% hydrogen peroxide (H_2_O_2_) in 40% methanol in TBS for 20 min. After washing, all the primary antibodies were detected by incubation with their corresponding secondary antibodies (1:400), Alex Fluor 488-conjugated goat anti-mouse IgG (H + L) (Molecular Probes, cat# A11029) and Alexa Fluor 568-conjugated goat anti-rabbit IgG (H + L) (Molecular Probes, cat# A11011), at RT for 2 hrs in a dark humidified environment. The stock solutions of primary and secondary antibodies were diluted in antibody dilution buffer (2% goat serum, 1% BSA and 0.025% Triton X-100 in TBS). After washing 5 times for 10 min with TBST, the slides were sealed with aqueous mounting medium, and confocal images were obtained with a Carl Zeiss LSM510 Meta microscope. For immunoperoxidase staining, permeabilized slice sections, prepared as described above, were incubated in 10 mM sodium citrate buffer with boiling for 5 min. Endogenous peroxidase was blocked using an endogenous peroxidase blocking buffer (3% H_2_O_2_ and 40% methanol in TBS), followed by 10% horse serum (Sigma Aldrich, cat# H1270) and 1% BSA with 0.025% Triton X-100. After washing, the sections were incubated with avidin and biotin (Vector Laboratories Inc., cat# SP-2001), followed by primary antibody at 4℃ for 16 hrs and secondary antibody for 2 hrs at room temperatures. The antibody-treated slides were incubated with ABC reagents (Vector Laboratories Inc., cat# PK-6100) for 30 min at room temperature. Finally, they were stained with DAB substrate (Vector Laboratories Inc., cat# SK-4105). The tissue integrity and neurodegeneration of the brains and eyes of *Drosophila* were visualized after hematoxylin and eosin staining (H&E staining). The H&E staining procedure was performed according to a previously described method^46^.

### Immunocytochemical confocal microscopic assay of mouse neurons

To observe the localization of α-Syn, HtrA2/Omi and mitochondria, primary neurons were isolated from the substantia nigra and striatum and cultured to 60% confluency using a standard primary cell culture method^47^. After staining with Mito Tracker Red CMXRos (Molecular Probes, cat# M7512) according to the manufacturer’s instructions, the neurons were fixed with 4% paraformaldehyde in PBS for 20 min and permeabilized with 0.5% Triton X-100 in TBS for 30 min at room temperature. The neurons were then blocked for immunohistochemical analysis. After washing with TBST, the neurons were treated overnight at 4℃ with a rabbit anti-α-Syn antibody (1:100) (Abcam, cat# Ab51252) or a rabbit anti-HtrA2/Omi antibody (1:100) (Abcam, cat# Ab64111). After washing, the neurons were treated with FITC-conjugated goat anti-rabbit IgG (H + L) antibody (1:200) (Abcam, cat# Ab6717) for 2 h at room temperature. To detect the localization of α-Syn, HtrA2/Omi and ER, neurons fixed with 4% paraformaldehyde in PBS were permeabilized and blocked as described above. After washing with TBST, the neurons were double-stained overnight at 4℃ with a rabbit anti-α-Syn antibody (1: 100) or a rabbit anti-HtrA2/Omi antibody (1:100) together with the ER marker mouse anti-PDI (1:100) antibody (Abcam, cat# Ab5484). After washing, the anti-α-Syn and anti-HtrA2/Omi antibodies were detected with secondary FITC-conjugated goat anti-rabbit IgG (H + L) antibody (1:400) (Abcam, cat# Ab6717), and the anti-PDI antibody was detected with Texas Red-conjugated goat anti mouse IgG (H + L) antibody (1:400) (Abcam, cat# Ab6787). The stock solutions of primary and secondary antibodies were diluted in antibody dilution buffer (2% goat serum, 1% BSA and 0.025% Triton X-100 in TBS). Confocal images were obtained with a Carl Zeiss LSM510 Meta microscope.

### Preparation of protein extracts from the substantia nigra and striatum

Four-week-old mnd2/mnd2, mnd2/+ and age-matched control mice were sacrificed to isolate the substantia nigra and striatum surgically. Total proteins were extracted from the substantia nigra and striatum of each mouse using a total protein extraction kit (Merck Millipore, cat# 2140) according to the manufacturer’s protocol. Total protein extracts of equal amounts from each sample were subjected to western blot analysis with a mouse anti-α-Syn antibody (Abcam, cat# Ab1903).

### Isolation of primary neurons from the substantia nigra and striatum

The substantia nigra and striatum were obtained surgically from four-week-old wild-type C57BL/6 mice to investigate the localization of α-Syn and HtrA2/Omi. The substantia nigra and striatum were minced into small pieces in dissection medium (DMEM/F-12, 32 mM glucose, 1% penicillin/streptomycin, and 0.5 mM L-glutamine), followed by the addition of 2 mL of HBSS and centrifugation at 300 × *g* for 5 min. The pellet was treated with 2 mg/mL papain in HBSS and 200 µg/mL DNase I solution for 30 min in a 30℃ water bath with a platform rotating at 150 rpm. The tissue was gently triturated by the addition of dissection medium to the single cell suspension and passage through a cell strainer (BD Falcon, cat# 08-771-19). The neuronal pellets were washed twice with ice-cold dissection medium by centrifugation at 300 × *g* for 7 min and resuspended in pre-warmed Neurobasal/B27 growth medium supplemented with 1% penicillin/streptomycin, 0.5 mM L-glutamine, 10 ng/mL FGF2 (Sigma Aldrich, cat# SRP4038), and 12.5 mM NaCl. The cells were seeded at a density of 120 cells/mm^2^ on poly-D-lysine and laminin-coated multiwell chambered cover slips (Grace Bio-Labs, Bend, Oregon) and incubated in a humidified incubator at 37℃ in an atmosphere of 5% CO_2_ and 95% air for 1 hr for neuronal attachment. After 1 hr, the spent medium was aspirated, followed by the addition of fresh growth medium and further incubation. Half of the spent medium was replaced every 3 days with the same volume of fresh pre-warmed growth medium containing 20 ng/mL FGF2.

### Preparation of *Drosophila* and mouse brain homogenates

Fly heads were homogenized using an ultrasonicator at 4℃ for 6 min at 40 kHz in 3 µL of total protein extraction kit TM buffer containing 0.1% SDS (Millipore, cat# 2140). The homogenized samples were centrifuged at 17,000 × g for 1 min at RT to remove the debris. The supernatant was collected and centrifuged again under the same conditions to collect the fresh supernatant. Four-week-old mnd2/mnd2 mice, mnd2/+ mice and aged match controls were sacrificed, and the brain tissues were chopped into small pieces, followed by the addition of 1 mL of total protein extraction kit TM buffer containing 0.1% SDS (Millipore, cat# 2140) to 0.2 g of brain samples. Samples were homogenized, and total proteins were isolated as described above.

### Pan-neuronal and eye-specific expression of transgenes in *Drosophila*

Pan-neuronal expression of transgenes was achieved by crossing virgin female flies (driver line) carrying elav-Gal4 on their X chromosome with transgenic flies. Eye-specific expression of transgenes was achieved by crossing virgin female flies (driver line) carrying GMR-GAL4 on their X chromosome with transgenic flies. These flies were maintained at 25℃ and, immediately after eclosion, sorted for western blotting, IHC, and survival and locomotion assays.

### Survival assay

Flies were maintained on standard cornmeal-sucrose-yeast-agar-molasses medium at 25℃ in a 60% humidified incubator (Han Baek Scientific Co., Seoul, South Korea) with a 12-hr light/dark cycle. Male transgenic flies were mated with virgin female elav-GAL4 driver flies. Newly eclosed flies were allowed to mature for 48 hrs, and then the male and female flies were separated into different jars. Exactly 100 adult female and 100 adult male flies were maintained for the aging experiments. During maintenance, the flies were transferred to fresh medium every 5 days, and their survival was recorded. This process was continued until all the flies had died. Non-age-related or non- disease-related death was censored. Analysis of the survival data was performed using the Kaplan-Meier method^48^.

### Locomotion assay

Transgenic flies were mated and maintained on standard cornmeal-sucrose-yeast-agar-molasses medium as described above for the locomotion assay. Fifteen flies were anesthetized with CO_2_ and placed in a 15-mL conical tube (SPL life sciences, cat# 51015) capped with cotton. Anesthetized flies were allowed to recover for 30 min at room temperature before the climbing assay. Flies were tapped to the bottom of the tube and allowed to climb with video recording for 30 sec. The experiments were repeated 3 times. After 10~15 sec, the numbers of flies remaining below the 2 mL mark (n_bottom_) and flies the 10 mL mark (n_top_) were recorded. The performance index (PI) was calculated for each group using the following formula: PI = 0.5x (n_total_ + n_top_ - n_bottom_)/n_total_, where n_total_ is the total number of flies, n_top_ is the total number of flies at the top, and n_bottom_ is the total number of flies at the bottom. If all flies climb to the top of the tube, the score is 1, and if no flies climb the score is 0^48^.

### Scanning electron microscopy study

Freshly sacrificed flies were dehydrated by serially transferring them into increasing ethanol concentrations of 30, 40, 50, 60, 70, 80, 90 and 100% for 10 min each at room temperature. The dehydrated flies were air dried prior to being preserved at -80℃. The dehydrated flies were mounted on a slide with one eye upward on black tape using colloidal graphite in an isopropanol base. The flies were fixed in osmium tetroxide and air dried prior to observation. All flies were placed on a rotating platform to permit orientation under a vacuum and imaged at 180 × magnification using a JSM-6400 scanning electron microscope (JEOL Ltd. Akishima, Tokyo, Japan).

### Statistics and reproducibility

All statistical analyses are reported as the mean ± SEM, and the significance was calculated using one-way ANOVA followed by Bonferroni/Tukey multiple tests for individual means using IBM SPSS statistics 21 software. *P* values less than 0.05 were considered statistically significant. When representative images are shown, at least three repeats were performed.

### Data availability

All data supporting the findings of this study are available from the corresponding author on request.

## ACKNOWLEDGMENTS

This research was supported by the Brain Research Program through the National Research Foundation of Korea (NRF) funded by the Ministry of Science, ICT & Future Planning (NRF-2017M3C7A1044818) and also partially funded by JINIS BDRD Research Institute.

## AUTHOR CONTRIBUTIONS

H.J.C. and M.A.M. J. designed the project; H.J.C., M.A.M.J. and M.M.R. designed the experiments; H.J.C., M.A.M.J., M.M.R., and H.J.K. analyzed results; H.J.C., M.A.M.J. and M.M.R. performed the experimental work; H.J.C., M.A.M.J., H. J.K. and S.T.H. wrote the manuscript. H. J.K. and S.T.H. supervised the project.

## ADDITIONAL INFORMATION

**Supplemental Information** accompanies this paper at https://doi.org/10.1038/ **Competing interest:** The authors declare no competing interests.

## Supplementary Information

### Supplementary Table

**Table S1.**
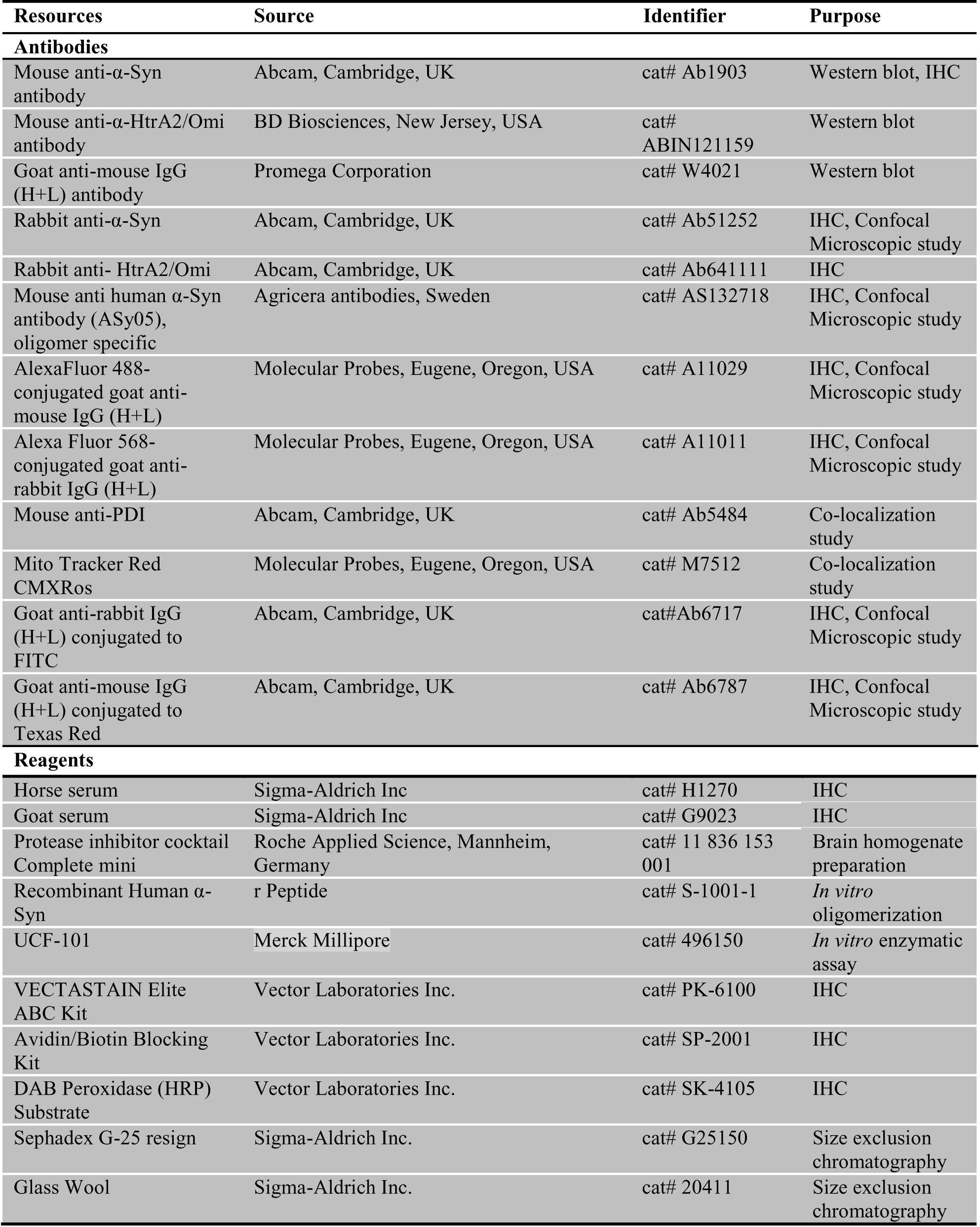
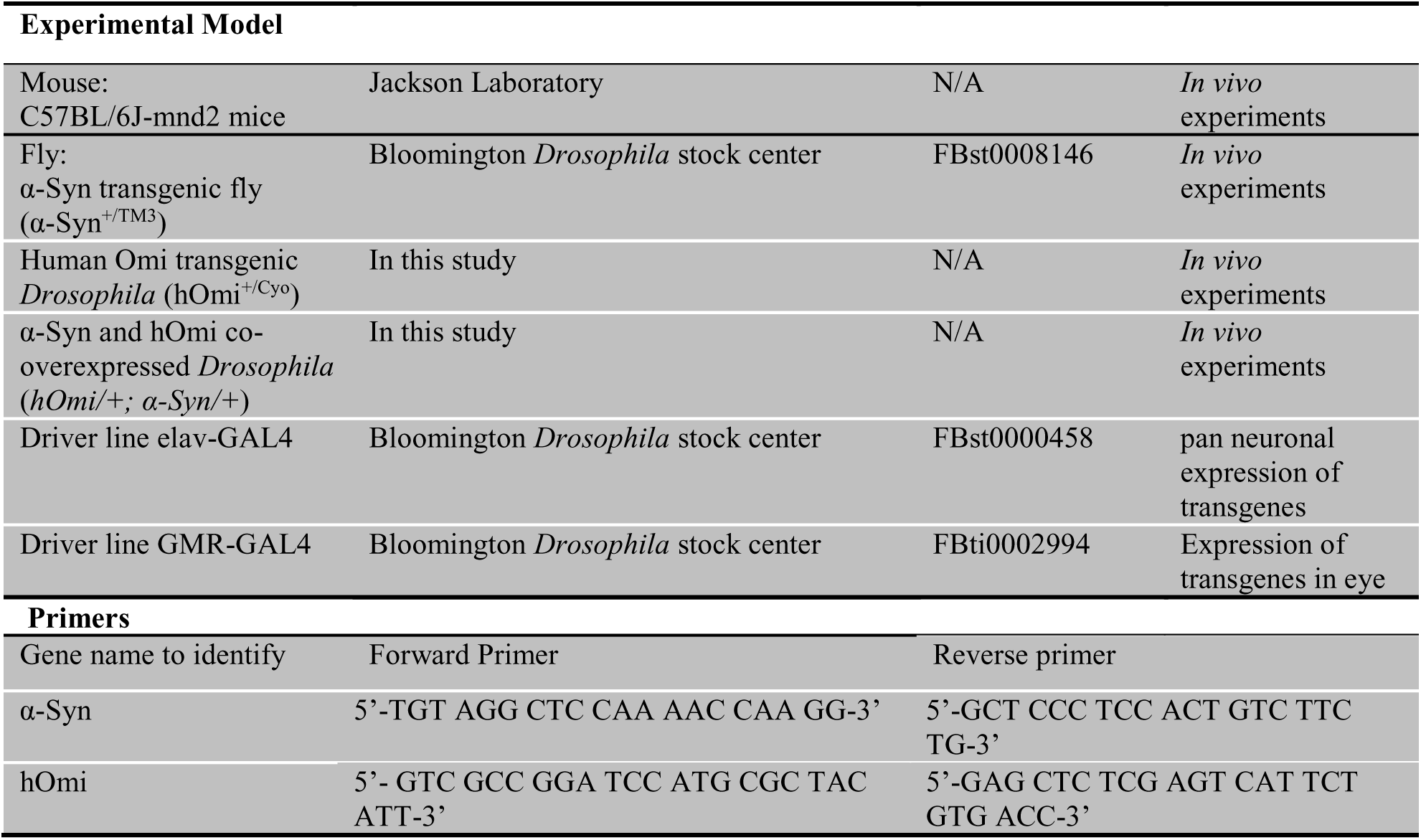
Resources used in experiments

### Supplementary Figures

**Figure S1.**
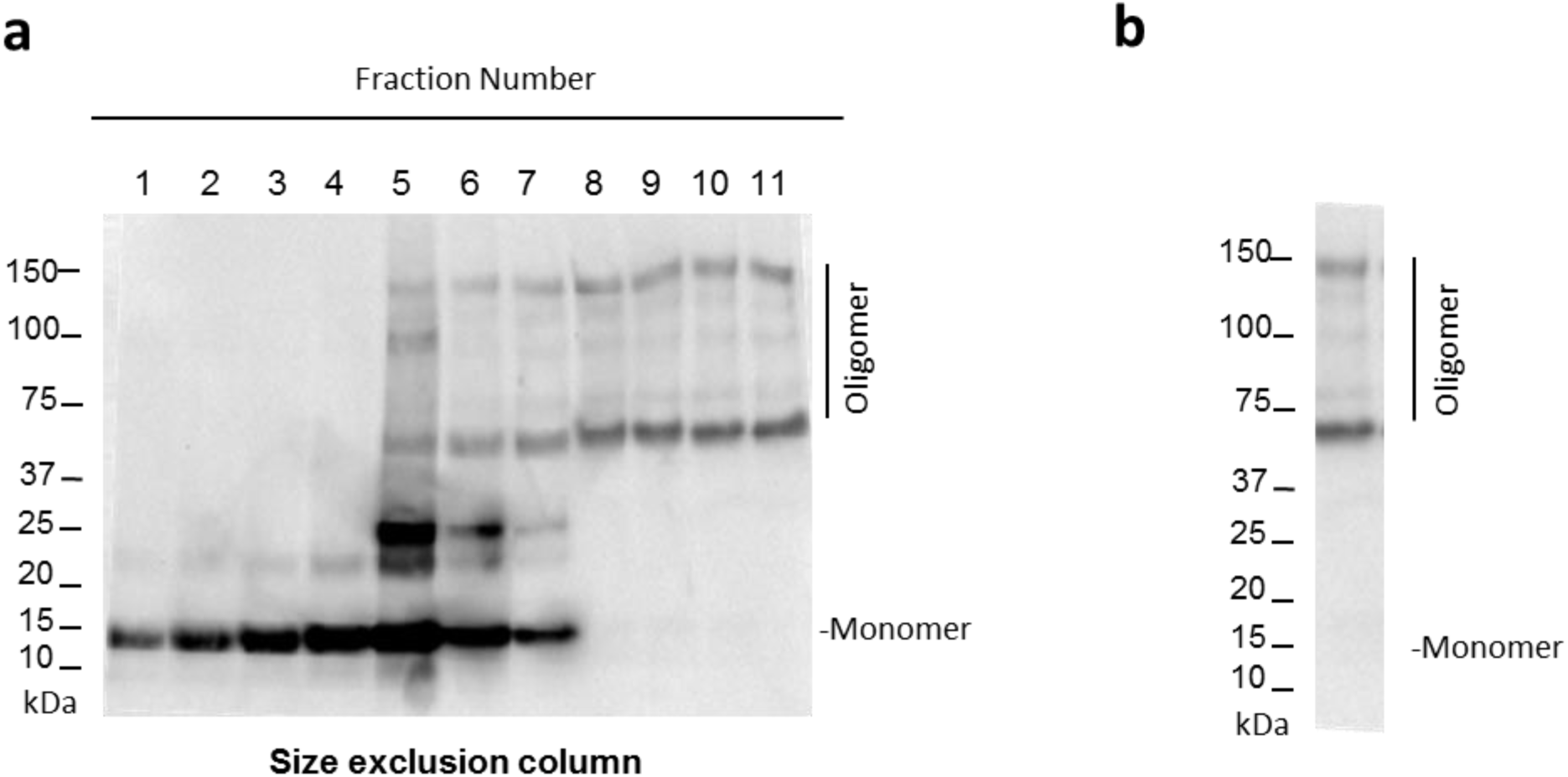
Oligomeric α-Syn was isolated using a size exclusion column. **a** SDS-PAGE gel showing oligomerized α-Syn fractionations after applying a size exclusion column. human recombinant monomeric α-Syn was oligomerized, and monomer, dimer, trimer, tetramer as well as higher order oligomers could be distinguished by molecular weight. **b** Isolated oligomeric α-Syn using the size exclusion column.

**Figure S2.**
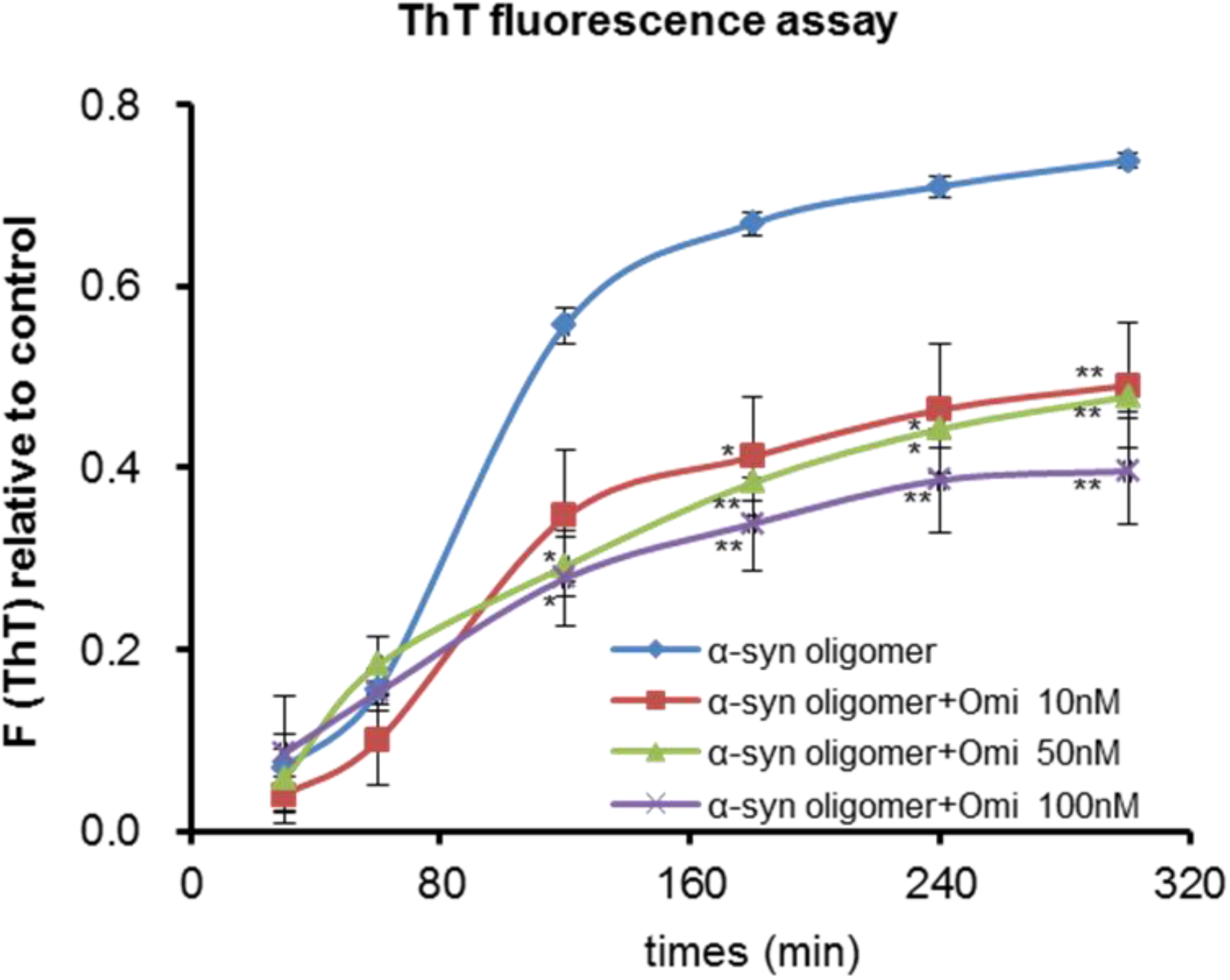
hOmi specifically degraded oligomeric α-Syn in a dose-dependent manner. The degradation of α-Syn was measured by the fluorescence intensity after staining of α-Syn with the oligomer-specific fluorescent dye ThT. Values represent the mean ± SEM from three independent experiments. ^*^*p*<0.05 and ^**^*p*<0.01.

**Figure S3.**
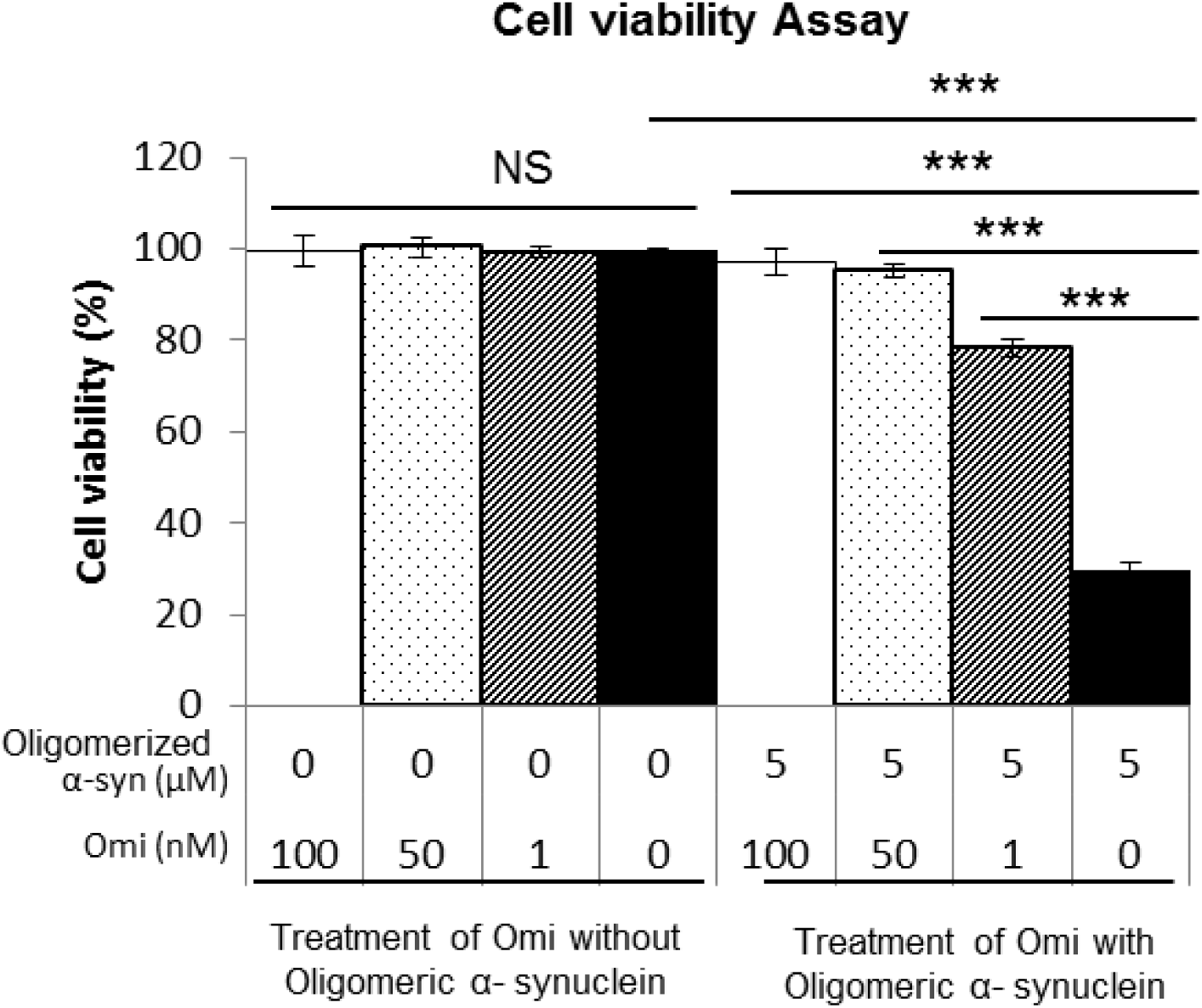
The neurotoxicity of oligomeric α-Syn was abolished after co-treatment with hOmi in a dose-dependent manner. Recombinant hOmi treatment was applied at 0 nM, 10 nM, 50 nM and 100 nM, and cell viability was assessed using the CCK-8 assay. Values represent the mean ± SEM from three independent experiments. NS, not significant and *^***^p*<0.001.

**Figure S4.**
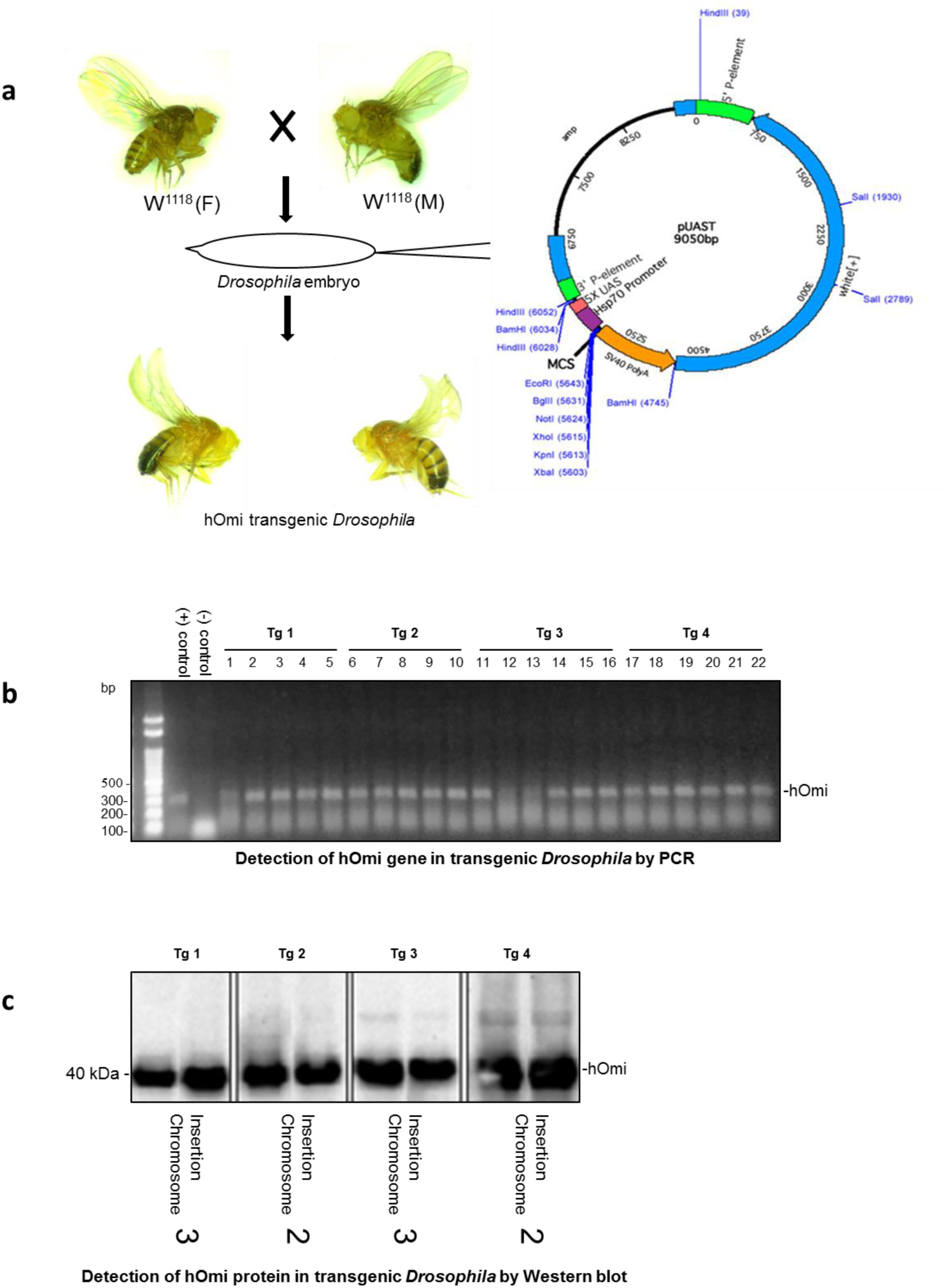
A transgenic *Drosophila* line expressing human HtrA2/Omi (hOmi) was successfully developed. **a** Schematic representation of the development of human HtrA2/Omi (hOmi) transgenic *Drosophila* lines. The full-length hOmi gene was cloned into the GAL4-responsive pUAST expression vector. Transgenic *Drosophila* lines of hOmi were generated by microinjection of a plasmid bearing the P element transposase under the control of the heat shock 70 (hs-π) promoter as a source of transposase into w^1118^ embryos. **b** Detection of the hOmi gene in 4 transgenic *Drosophila* lines by PCR. **c** Detection of the hOmi protein in 4 transgenic *Drosophila* head homogenates by western blotting, in which head homogenates were obtained from the progeny of hOmi crossed with elav-GAL4 flies.

**Figure S5.**
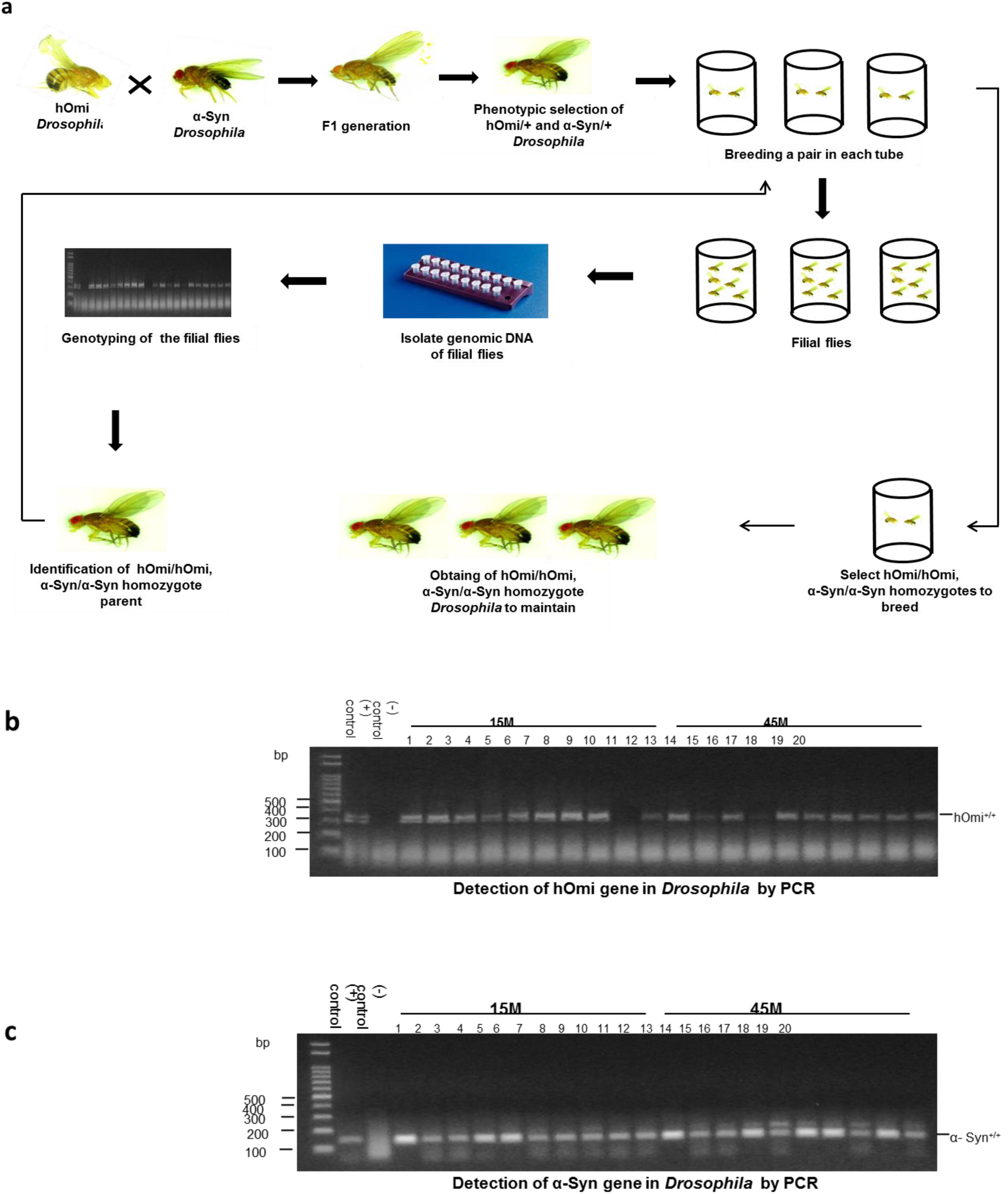
A Transgenic *Drosophila* line carrying both hOmi and α-Syn genes as homozygotes was developed by massive PCR screening. **a** Schematic representation of developing *Drosophila* lines carrying both hOmi and α-Syn genes as homozygotes by crossing hOmi transgenic *Drosophila* Tg4 (X/Y; hOmi/Cyo; +/+) with α-Syn transgenic *Drosophila* (X/Y; +/+; α-Syn/α-Syn). **b** Example of 2 transgenic *Drosophila* lines carrying hOmi as homozygotes. The *Drosophila* lines carrying hOmi as homozygotes were assessed by genotyping fly progenies. If all the progenies from either parent crossed with control carried hOmi, the parental flies were identified as homozygotes and used in this work. **c** Example of 2 transgenic *Drosophila* lines having α-Syn as homozygotes. The *Drosophila* lines having α-Syn as homozygotes were assessed by genotyping the fly progenies. If all the progenies from either parent crossed with the control had hOmi, the parental flies were identified as homozygotes and used in this work.

**Figure S6.**
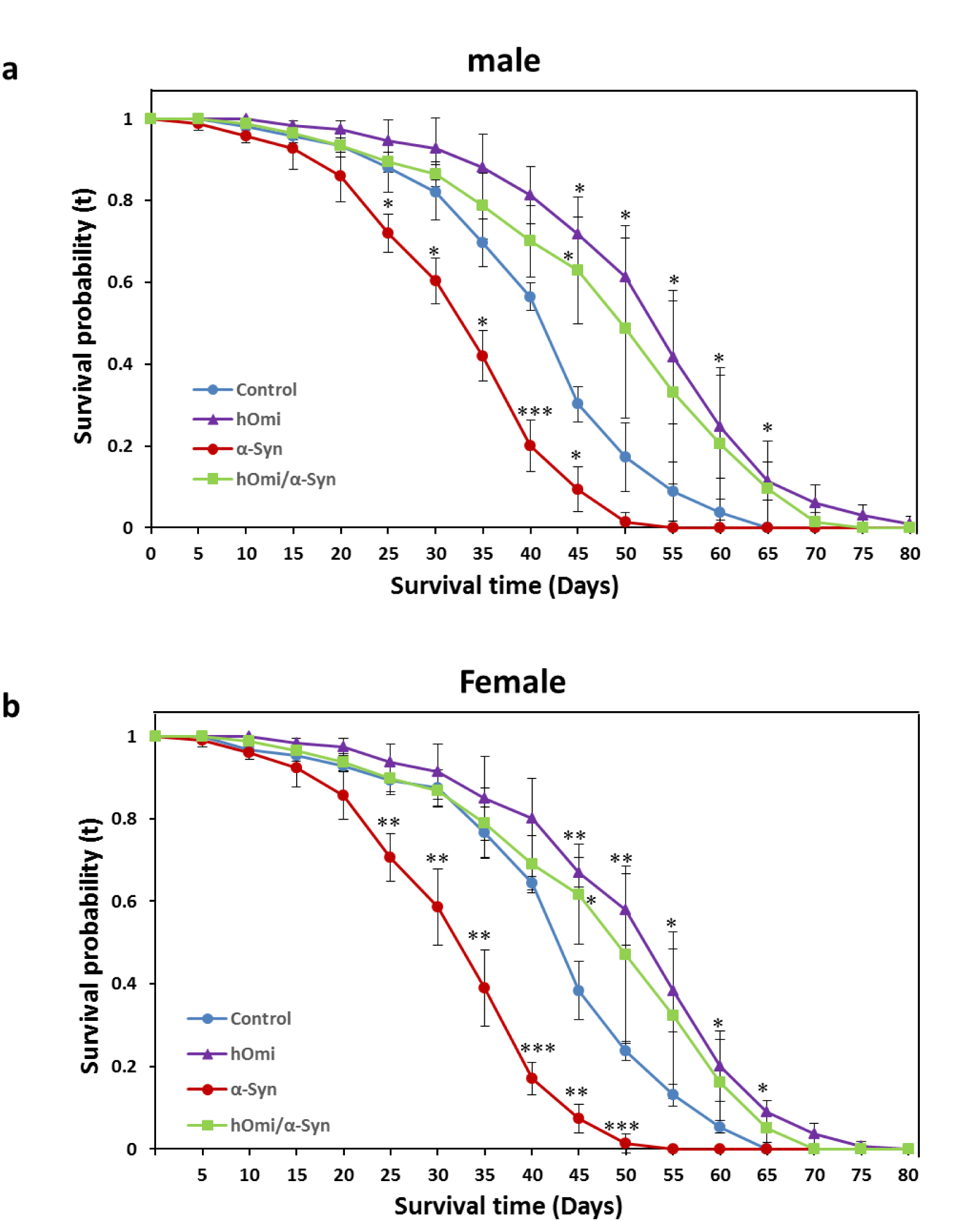
The Kaplan–Meier Survival assay measured by the survival rate of hOmi, α-Syn, or hOmi/α-Syn flies demonstrated a neuroprotective role of hOmi against α-Syn-induced cytotoxicity in both male a and female b flies. Values are the mean ± SEM from three independent experiments. NS, not significant, ^*^*p*<0.05, *\*\*p*<0.01, ^***^*p*<0.001.

